# A comprehensive map of the ageing blood methylome

**DOI:** 10.1101/2023.12.20.572666

**Authors:** Kirsten Seale, Andrew Teschendorff, Alexander P Reiner, Sarah Voisin, Nir Eynon

**Author notes:** Corresponding Author: Professor Nir Eynon, Australian Regenerative Medicine Institute, Monash University Clayton, VIC 3800, Australia. Sarah Voisin & Nir Eynon are co-senior authors.

## Abstract

During ageing, the human methylome exhibits both differential (i.e. change in mean) and variable (i.e. change in variance) shifts, along with a general rise in entropy. However, it remains unclear whether DNA methylation sites that increasingly diverge between people (i.e. variably methylated positions (VMPs)) are distinct from those undergoing changes in mean methylation levels (i.e. differentially methylated positions (DMPs)), which changes drive entropy, how they contribute to epigenetic age measured by epigenetic clocks, and whether cell type heterogeneity plays a role in these alterations. To address these questions, we conducted a comprehensive analysis using > 32,000 human blood methylomes from 56 datasets (age range = 6-101 years). Our findings revealed an unprecedented proportion of the blood methylome that is differentially methylated with age (48% DMPs; FDR< 0.005) and variably methylated with age (37% VMPs; FDR< 0.005), with many sites overlapping between the two groups (59% of DMPs are VMPs). We observed that bivalent and Polycomb regions become increasingly methylated and divergent between individuals, while quiescent regions lose methylation in a more homogeneous manner between individuals. Unexpectedly, both chronological and biological clocks, but not pace-of-aging clocks, show a strong enrichment for those CpGs that accrue both mean and variance changes during aging. Furthermore, we uncovered that it is the accumulation of DMPs shifting towards a methylation fraction of 50% that drive the increase in entropy, resulting in an overall smoothening of the epigenetic landscape. However, approximately a quarter of DMPs oppose this direction of change, exhibiting anti-entropic effects. While DMPs were mostly unaffected by changes in cell type composition, VMPs and entropy measurements showed moderate sensitivity to such alterations. This investigation represents the largest to date of genome-wide DNA methylation changes and ageing in a single tissue, offering valuable insights into primary molecular changes that hold meaning for chronological and biological ageing.

## Introduction

All humans experience similar ageing symptoms with chronological time, however, the degree and speed at which these changes occur varies between individuals, leading to inter-individual differences in the time of onset and severity of age-associated disease and disability (i.e. individuals who are the same chronological age will differ in their ‘biological’ age)^1^. Ageing is initiated at the basic level of biological organisation and is governed by 12 interconnected hallmarks^2^. Alterations to the epigenome are a primary hallmark of ageing, and a focus of this research^2^. Specifically, the age-associated changes in DNA methylation (DNAm), which accrues numerous, widespread changes. Of importance are two distinct linear changes: *differential* and *variable* patterns of DNAm. Age-associated *differentially methylated positions* (DMPs) capture age-related DNAm changes that are *shared* between individuals, whereas age-associated *variably methylated positions* (VMPs) capture DNAm changes that *diverge* between people over the lifespan^3^. It is therefore plausible that at the epigenetic level, two individuals with identical chronological ages (and patterns of DMPs) may display divergent patterns across VMPs. Albeit DMPs and VMPs are not mutually exclusive (i.e. a CpG can be both a DMP and a VMP), both features contribute to age-associated DNAm changes, but in fundamentally different ways. Insights into epigenetic ageing can also be drawn from quantifying the changes at the whole methylome level using single measurements of *entropy;* a probability formula that estimates the amount of information in a set of CpGs averaged across a population of cells^3^. In blood, an increase in entropy over time implies increasing age-related ‘methylation disorder,’ implying the methylome loses information with age^3^. Entropy can summarise all the age-related changes in DNAm using a single value, for a sample at a particular age, making it a highly useful in capturing the entirety of the age-related changes in the methylome. Previous literature has shown that DMPs shifting towards a methylation fraction of 50% drive entropy^4,5^, but there are also subsets of DMPs that trend away from the mean and may oppose or counteract the effect^6^.

The vast majority of studies have solely investigated DMPs, including epigenetic clocks, which may favour DMPs for accuracy of prediction. However, focusing only on patterns of DMPs is limiting when trying to understand aspects of biological ageing, particularly when making sense of why individuals of the same age display vastly different ageing rates. Yet our knowledge of the extent to which the methylome is differentially and variably methylated with age is limited, since much of our understanding to date has been drawn from smaller, isolated datasets, such as Slieker et al. (∼ 3,000 samples) and Hannum et al. (∼650 samples)^4,7^. Another complexity in human studies is that cohorts are also highly heterogenous in their characteristics (e.g. sex distribution, disease status, ethnicity, age), making it challenging to generalise findings drawn from individual cohorts. Furthermore, detecting DMPs and especially VMPs, requires a large sample size and a broad age range, so it is possible that important epigenetic ageing patterns have been missed in previous studies due to insufficient statistical power. Moreover, no studies to date have investigated the contribution of VMPs to changes in entropy.

To address these gaps, we performed a large-scale epigenome-wide association study (EWAS) meta-analysis of age in whole blood. Leveraging the statistical power from 56 whole blood datasets of > 32000 samples, we quantified the age-associated DNAm changes, distinguishing between *shared* changes (DMPs) and *divergent* changes (VMPs) over the life course. In addition, we sought to understand how entropy changes and how DMPs and VMPs contribute to these entropy dynamics, including those linear changes that trend towards and away from the mean. We also explored the effect of cellular heterogeneity on these age-associated signatures, since bulk tissue analyses reflects the aggregate cell type DNAm changes as well as age-related changes in cellular composition. We performed cell type correction in addition to an unadjusted analysis^8^.

With unprecedented statistical power and a comprehensive investigation of epigenetic signatures of ageing in blood, our study shows the sheer scale of differential and variable methylation in human ageing, with important implications for chronological and biological ageing. Such insights will serve as a foundation for studying the effects of lifestyle, dietary, or pharmaceutical interventions on ageing signatures, an approach that was recently shown by our group to successfully capture the rejuvenating effects of exercise training on age-associated DMPs in muscle^9^.

## Results

### Methodology overview

Briefly, the first step in the methodology was to gather existing DNAm datasets from public open access and controlled access data repositories, to assemble an exhaustive database of DNAm profiles in blood (**Figure 1, Extended Data Fig. 1**). We collected 56 whole blood datasets with a combined sample size of 32,136 samples (**Extended Data Fig. 1**). This large database of human methylomes spanned a broad age range (6 – 101 years) and laid a solid foundation to quantify DMPs, VMPs, and entropy in each dataset.

**Figure 1.**
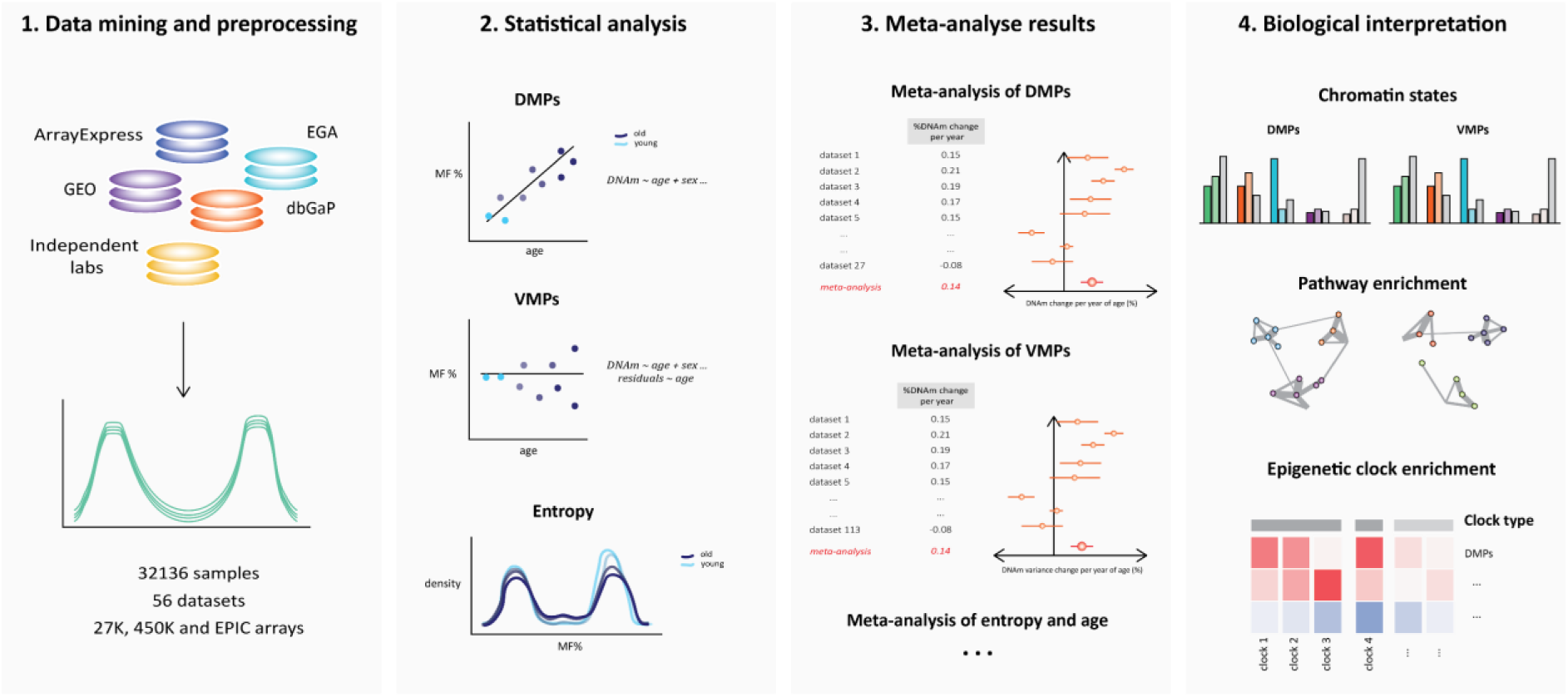
Overview of methodology. Raw DNAm profiles from the 27K, 450K and EPIC Illumina Array platforms were sourced from open-access and controlled access databases, including ArrayExpress, GEO, EGA, dbGaP and independent labs. The DNAm profiles of 32,136 samples were collected and pre-processed, for both males and females, from 56 datasets in whole blood (top panel). The relationship between age and average DNAm, or the relationship between age and DNAm variance, or the relationship between age and entropy was estimated in each dataset independently (second panel). For each CpG and for entropy, summary statistics were then meta-analysed across datasets to identify DMPs, VMPs and to determine whether entropy was significantly associated with age. We performed a meta-analysis of age for DMPs, VMPs and entropy, by pooling the results from the independent EWAS (third panel). We then performed a functional analysis of the DMPs and VMPs to interpret the findings, including pathway analysis, and enrichment in chromatin states (fourth panel).

Our analysis began with identifying the linear age-related DNAm changes at individual CpGs (i.e. DMPs and VMPs). To identify DMPs, we completed an independent EWAS by fitting a linear model for each dataset, regressing DNAm against age and other covariates that are known to modulate DNAm levels (i.e. sex, ethnicity, batch, body mass index (BMI)) (**Supplementary Table 1**). We then extracted the summary statistics (i.e. the age-related change in DNAm level) from each EWAS and conducted an inverse variance based fixed-effects *meta-analysis of differential methylation and age*. To identify VMPs, we followed a similar approach, using the Breusch-Pagan test for heteroscedasticity in each independent dataset, which models the change in DNAm variance as a function of age. The summary statistics extracted from the independent EWAS (i.e. the age-related change in DNAm variance) were then pooled using a sample-size based fixed effects *meta-analysis of variable methylation and age*. We then compared age-related CpGs that were only DMPs (*homoscedastic DMPs*), those that were only VMPs (*constant VMPs*) and those that were both DMPs and VMPs (*DMPs-VMPs*). Finally, we compared DMPs that carried different degrees of information, as defined by Shannon entropy: we compared DMPs that trend *towards* the mean (*entropic* DMPs) with those that trend *away* from the mean towards fully methylated and unmethylated states (*anti-entropic* DMPs).

Finally, we performed a comprehensive entropy analysis by looking at *genome-wide Shannon entropy*. We took the same statistical approach as with the DMP and VMP analyses described above: first, we estimated the strength of the association between age and Shannon entropy in each independent cohort; then, we pooled these effect sizes across the different cohorts using a fixed-effects meta-analysis to obtain an overall meta-analysis effect size of *change in entropy per decade of age* (**Supplementary Table 2**). We also calculated entropy on the *age-related CpGs* (i.e. the complete list of DMPs and VMPs) and the remaining non-age-related CpGs, and meta-analysed the results to compare the *change in entropy with age at age-related CpGs vs non-age-related CpGs,* evaluating whether it is these age-related shifts driving changes in entropy. Then, we took a more granular approach and investigated whether different classes of DMPs and VMPs contribute differently to entropy measurements (homo-DMPs, constant VMPs, DMPs-VMPs, or entropic vs anti-entropic DMPs). This allowed us to confirm whether it is the *differential* or *variable* shifts in DNAm that increase entropy, or *both*.

Throughout the pipeline, we compared these different classes of age-related changes in terms of genomic location (chromatin states profiled in PBMCs^10^, annotated thank to <u>a comprehensive annotation of the Illumina</u> <u>Methylation arrays</u>^11^), biological pathways (*<u>gene ontology</u>* (GO) terms, *<u>human phenotype ontology</u>* (HPO) terms, *<u>canonical pathways</u>* (CP), *<u>expression signatures of genetic and chemical perturbations</u>* (CGP), and *<u>immunologic</u> <u>signatures</u>* (C7)), and overrepresentation in various epigenetic clocks (chronological age: Horvath’s pan-tissue clock^12^, Hannum’s clock^4^, the blood clock developed by Zhang et al. 2019^13^, the centenarian clock^14^, and the mammalian universal clock^15^; biological age: PhenoAge^16^; pace of aging: DunedinPoAm^17^, DunedinPACE^18^). We also explored the effect of cellular heterogeneity on these age-associated signatures. CpGs that determine cell identity are typically lowly methylated in a given cell type, while being highly methylated in other cell types, and the overall methylation fraction in bulk tissue at those cell-type-specific CpGs would be highly sensitive to changes in the relative proportions of different cell types. Ageing is associated with an increase in monocytes, neutrophils, basophils, NK cells, CD4+ and CD8+ T memory cells, with a concomitant decrease in naïve B cells, T-regulatory cells, CD4+ and CD8+ naïve T cells^19^. We deconvoluted the proportions for granulocytes, monocytes, Natural Killer cells (NK), CD4+ T cells, CD8+ T cells and B cells for each sample using a reference-based method^20^, and repeated all the above-mentioned analyses after adjusting the linear model for blood cell type proportions.

### Ageing is associated with widespread changes in DNAm levels and increases in DNAm variance in blood

With the unprecedented statistical power granted by > 32,000 samples from 56 datasets, we found that nearly half of all tested sites (333,300 CpGs) were DMPs (48%) at a stringent FDR < 0.005. Two-thirds of DMPs (66%) decrease in DNAm levels (‘hypoDMPs’), while the remaining third increase in DNAm levels with age (‘hyperDMPs’) (**Figure 2A**). HyperDMPs increase by an average of 0.027% methylation fraction per year of age, noting a maximum increase of 0.46% per year of age for cg26079664, and hypoDMPs decrease by an average of -0.034% per year of age, with the maximum decrease of -0.55% per year of age for cg10501210. Our meta-analysis identified DMPs that were highly consistent across datasets. For example, cg16867657, which is in the promoter of *ELOVL2* and has been associated with ageing in a plethora of studies^21–24^, was estimated to gain 0.45% DNAm per year of age across the different datasets (**Figure 2B**).

**Figure 2.**
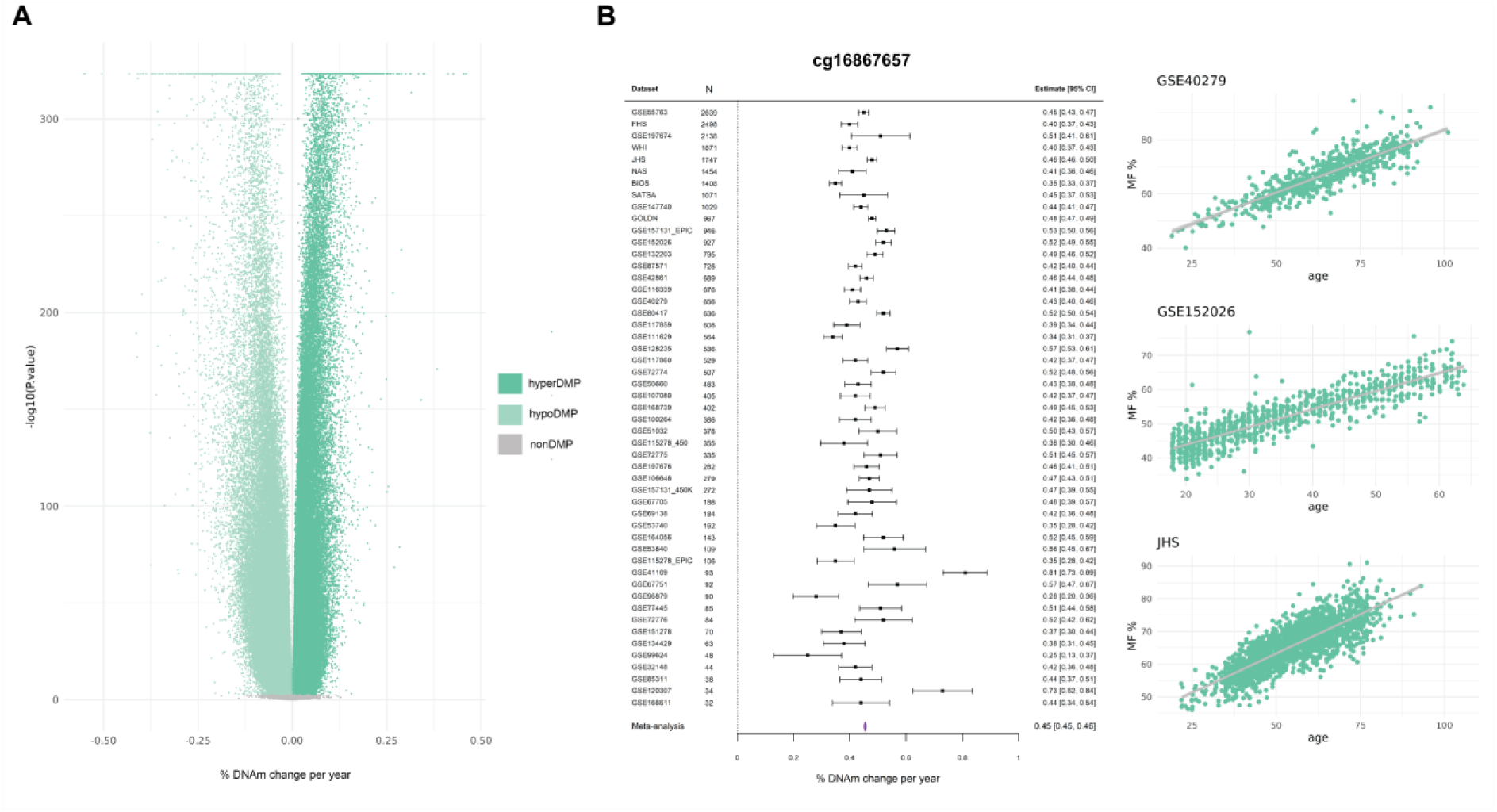
Meta-analyses of EWAS in blood to identify DMPs. **A)** A volcano plot displaying the meta-analysis effect size (x-axis) and significance (y-axis) for the 696,228 tested CpGs in the differential methylation meta-analysis of age in blood. The strongest associations have the smallest *p*-values and will be the highest points on the plot. Hypomethylated (hypoDMP), hypermethylated (hyperDMP) and non-DMPs are represented by the colours in the legend. DMPs are classified at a false discovery rate (FDR) < 0.005. The ‘ceiling’ of extremely significant p-values is an artefact of the software that cannot handle numbers smaller than 2.2 x 10^-16^. **B)** A forest plot of a highly significant CpG (cg16867657). The dataset and sample size are on the left side of the plot, with the corresponding effect size and errors represented by the point and error bars. The meta-analysis effect size is represented by the purple polygon. The methylation plots for this CpG from three independent blood cohorts, including GSE40279, GSE152026 and Jackson Heart Study (JHS) are displayed on the far right, with age on the x-axis and methylation fraction (MF) as a percentage on the y-axis. Each point on the plot represents a single sample.

There was an inverse correlation between the overall methylation fraction of a CpG and the direction of change during ageing: DMPs whose DNAm levels were usually high (> 75% on average), were overwhelmingly hypoDMPs, while DMPs whose DNAm levels were usually low (< 25% on average), were overwhelmingly hyperDMPs (**Extended Data Fig. 2**). In contrast, DMPs with intermediate DNAm levels trend equally frequently towards high and low DNAm levels. For example, in the BIOS dataset (n = 1408), 21% of DMPs were considered to have an ‘intermediate’ methylation level, of which 40% gained methylation with age, and 60% lost methylation with age (**Extended Data Fig. 2**).

HypoDMPs were over-represented in quiescent chromatin regions and those weakly repressed by Polycomb complexes, while hyperDMPs were over-represented in bivalent promoters and enhancers as well as regions repressed by Polycomb complexes (ꭓ^2^ test *p-*value < 2.2e-16) (**Extended Data Fig. 3A**). Despite being located in distinct chromatin states, hypoDMPs and hyperDMPs were found in similar genes (Fisher’s exact test p-value < 2.2e-16), related to e.g. signal transduction & signaling (GO), developmental conditions (HPO), naïve to memory T-cell (MSigDB immunologic gene set) (**Extended Data Fig. 3B**). With the exception of the universal pan-mammalian clock, all chronological and biological clocks were enriched for both hypo and hyperDMPs, with no difference in enrichment for these two classes of DMPs in biological vs chronological clocks (Fisher’s exact test FDR < 0.005, **Extended Data Fig. 3C).** Pace of aging clocks did not show any enrichment for DMPs (Fisher’s exact test FDR > 0.005, **Extended Data Fig. 3C).**

These results remain largely unchanged when the meta-analysis was adjusted for cellular heterogeneity (Pearson’s correlation of meta-analyses effect sizes = 0.94, *p* value < 2.2e-16) (**Extended Data Fig. 4A**).

We then meta-analysed the same 56 whole blood datasets to identify changes in methylation variability (VMPs) during ageing. We identified 243,958 VMPs (37% of tested CpGs) at FDR < 0.005, nearly all of which *increased* in variance (99% of VMPs). The magnitude of the age-related changes in variance is small, for example, the average increase in variance across all datasets for the most significant VMP, cg21899500, is 0.01% per year of age (**Figure 3A**). There was a large overlap between DMPs and VMPs (i.e. a CpG site whose average DNAm level changed during ageing was also more likely to see its variance increase with age; Fischer’s exact test *p* value < 2.2 x 10^-16^). We identified 196,192 DMPs-VMPs, 137,108 homoscedastic DMPs (i.e. DMPs only), and 47,766 constant VMPs (i.e. VMPs only) (**Figure 3B**). Among the DMP-VMPs, 73,357 (37%) increased in both average methylation and variance, and 122,835 (63%) decreased in average methylation but increased in variance.

**Figure 3.**
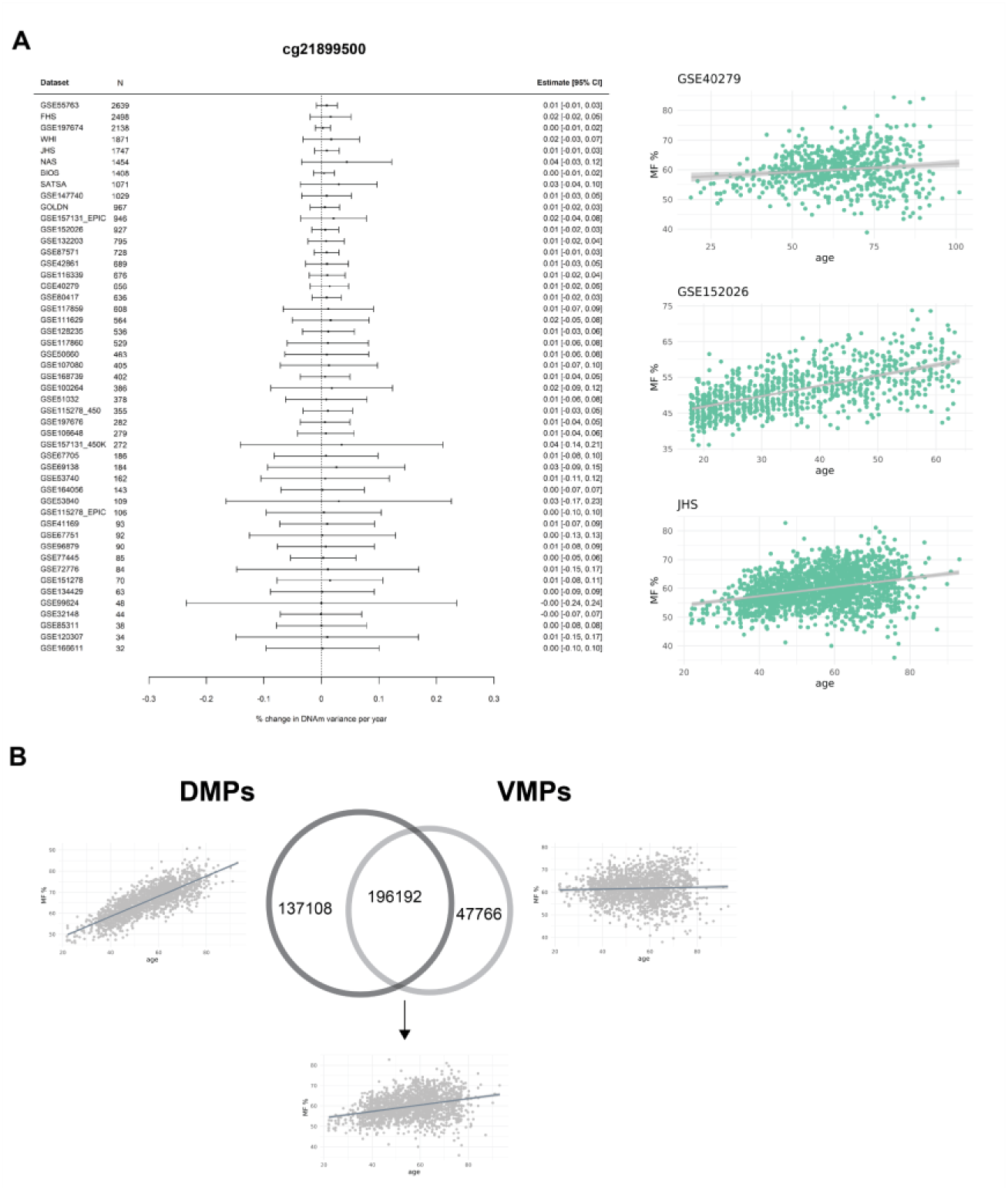
Meta-analyses of EWAS in blood to identify VMPs. **A)** A forest plot of the top VMP (cg21899500) that is hypermethylated with age and increases in variance with age, and the three adjacent methylation plots from three independent datasets, GSE40279, GSE152026 and JHS. **B)** Venn diagram of the overlap between DMPs and VMPs.

We compared the distributions of homoscedastic DMPs, constant VMPs, and DMPs-VMPs in different chromatin states profiled in PBMCs^10^. Constant VMPs were enriched in enhancers and Polycomb regions, DMPs-VMPs in bivalent promoters and enhancers as well as Polycomb regions, and homoscedastic DMPs were in quiescent regions (ꭓ^2^ test *p-*value < 2.2e-16) (**Extended Data Fig. 5**Error**! Reference source not found.A**). The three classes of age-related changes were related to a plethora of pathways, very similar to hypo and hyperDMPs (**Extended Data Fig. 5B**). All chronological and biological clocks were strongly enriched for DMPs-VMPs, while pace of aging clocks were not enriched for any kind of age-related CpGs (**Extended Data Fig. 5C).** Homoscedastic DMPs were overrepresented in the pan-tissue clock but depleted in Zhang et al.’s clock. Constant VMPs were depleted in two chronological clocks but were overrepresented in the PhenoAge.

We repeated the VMP meta-analysis after adjusting for blood cell type composition and as for the DMP analysis, results remained largely unchanged (Pearson’s correlation of meta-analyses Zscore = 0.93, *p-*value < 2.2e-16) (**Extended Data Fig. 4B**). However, VMPs seemed to be more sensitive than DMPs to confounding by cell type proportions, as more than a third of VMPs (37%) were *only* significant in the meta-analysis **not** adjusted for cell types. With that said, an additional 5,913 CpGs were classified as VMPs (4%) only after we adjusted for cell types. We identified 159,166 VMPs (22% of tested CpGs) after correcting for cell type composition.

### Entropy increases in the ageing blood methylome, driven by the cumulative changes in differential but not variable methylation at entropic CpGs

We determined whether the ageing blood methylome increases in entropy (‘chaos’) with age, and what type of epigenetic changes underpin this phenomenon. Entropy captures the amount of information encoded by the epigenome: if a CpG is highly (∼100%) or lowly (∼0%) methylated, this implies that said CpGs is highly “predictable” over all cells in a given sample; conversely, if a CpG has a methylation fraction closer to 50%, it is deemed “unpredictable” across cells within a sample. As the methylation state of genes determines cellular identity and therefore cellular function, entropy (i.e. ‘chaos’) increases when multiple CpGs throughout the genome drift towards a methylation fraction of 50%. An entropy of 0 means that every CpG is either methylated at 0% or 100%, and an entropy of 1 means that every CpG is methylated at exactly 50%^3^. In these two opposite scenarios, the methylome of a cell is either entirely predictable, or entirely unpredictable.

When taking *all* CpGs into account (both age- and non-age-related CpGs), we observed a very small but significant increase in entropy of 0.0005 per decade of age (*p-*value < 0.0001), with substantial heterogeneity between cohorts (*I*^2^ = 88%) (**Extended Data Fig. 6**).

As an increase in entropy with age reflects a drift towards a methylation fraction of 50% over multiple CpGs, we hypothesised that the increase in entropy would be driven by age-related CpGs (DMPs and/or VMPs). We re-calculated entropy in each sample from each dataset, but only taking into account the methylation levels at age-related CpGs (Error! Reference source not found.). As a ‘control’, we also re-calculated entropy in each sample from each dataset, but only taking into account the methylation levels of non-age-related CpGs. In line with our hypothesis, we found that non-age-related CpGs do *not* contribute to the global increase in entropy with age, with a meta-analysis effect size of -0.0003 change in entropy per 10 years of age (*p*-value <0.0001, *I*^2^ statistic 46%) (**Figure 4A**). In contrast, age-related CpGs increase in entropy by 0.002 per decade of age (*p*-value <0.0001, *I*^2^ statistic 85%) (**Figure 4A**). Moreover, the baseline entropy (i.e. the entropy value at the youngest age in a particular dataset) for the non-age-related sites is lower than the baseline entropy for the age-related sites (**Figure 4B**). This can be explained by the fact that there are more CpGs with intermediate methylation levels among age-related sites (∼20%), than non-age-related sites (∼1%).

**Figure 4.**
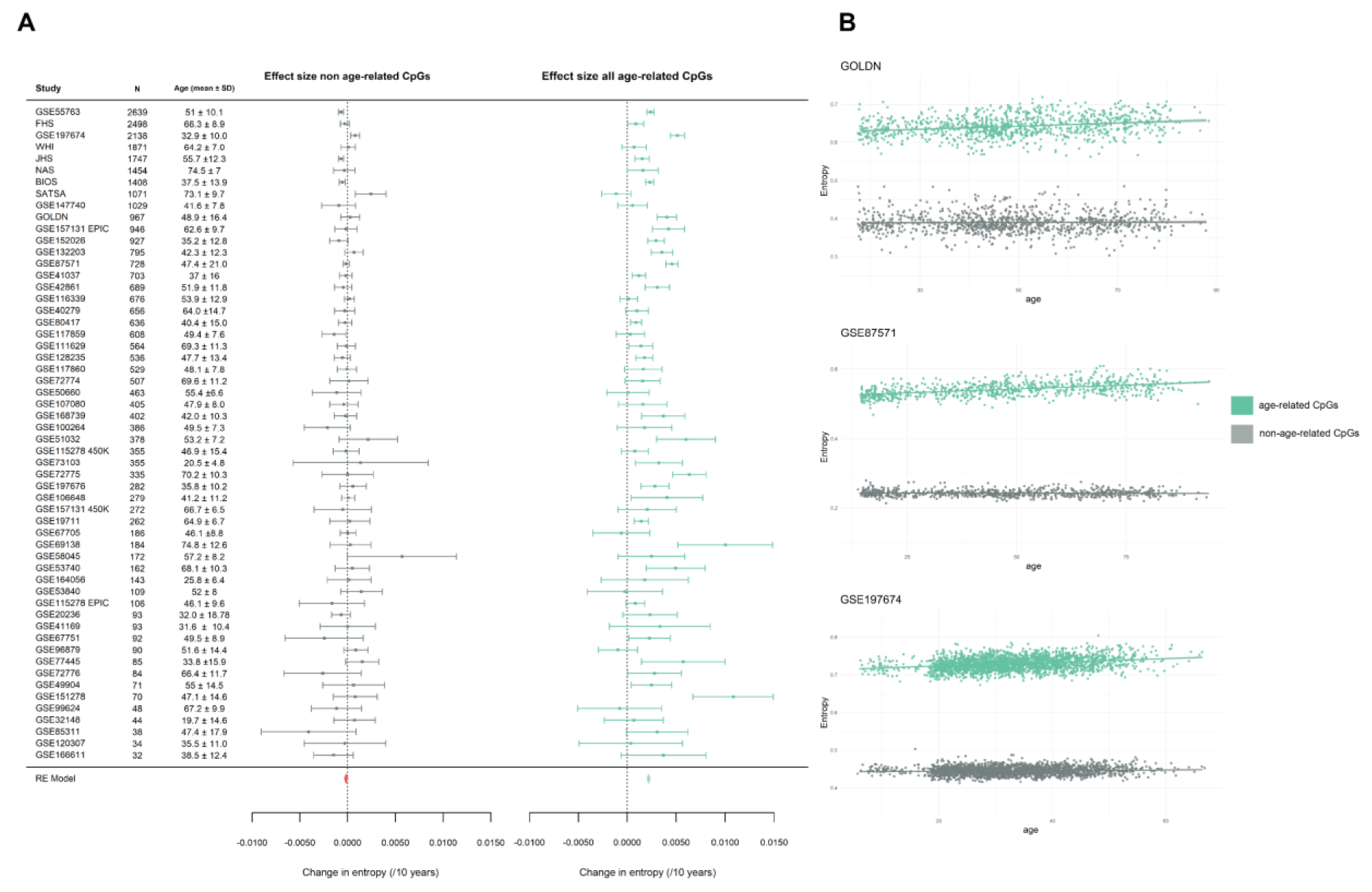
Meta-analysis of entropy and age. **A)** A forest plot of the two meta-analyses comparing the changes in entropy between the non-age-related CpGs (left) and the age-related CpGs (right) identified in blood. The meta-analyses effect sizes are represented by the orange and light green polygons. On the x-axes is the change in entropy per decade of age, and on the y-axes are the effect sizes and standard error measurements from the independent EWAS. The dataset name, sample size (N) and age ± sd (standard deviation) are to the left of the forest plot. **B)** Three graphs with entropy (y-axis) plotted against age (x-axis) for the age-related CpGs and non-age-related CpGs from three independent blood datasets, GOLDN, GSE87571 and GSE197674.

To dissect the respective contributions of differential or variable methylation to changes in entropy, we calculated entropy for two datasets with a large sample size and broad age range (BIOS and FHS) (**Supplementary Table 1**) on homoscedastic DMPs, DMPs-VMPs, and constant VMPs (**Error! Reference source not found.A**, S**upplementary Table 3**), and regressed entropy against age in each category. DMPs-VMPs display the largest significant increase in entropy during ageing. We also observed that entropy significantly increases at homoscedastic DMPs, however, is lower both overall and at baseline than for DMPs-VMPs, which reflects the *type* of CpG affected by differential methylation or by a change in variance. While both DMPs and VMPs affect CpGs whose methylation levels start at high or low levels, VMPs have a greater proportion of CpGs with *intermediate* DNAm levels at baseline (∼28% of VMPs are intermediately methylated vs ∼20% of DMPs). While the overall entropy at CpGs that are only VMPs (i.e. constant VMPs) is high, we observed a decrease in entropy at those sites that was significant in only one of the two examined datasets (**Error! Reference source not found.A, Supplementary Table 3**C), suggesting that it is the differential shifts in DNAm towards the mean that contribute to the overall increases in entropy with age.

To further our investigation into the contribution of DMPs to changes in entropy, we used the BIOS blood dataset, which has a large sample size and distribution of samples across a large age range (**Extended Data Fig. 1**, **Supplementary Table 1**). Although the majority of DMPs (∼73%) *converge* to the mean with age, one third of DMPs (∼27%) *diverge* away from the mean towards high and low methylation fractions (**Figure 5A**). To determine this effect on entropy, we then recalculated entropy on the converging and diverging DMPs, respectively. Remarkably, we found a highly significant *increase* in entropy in the converging sites of 0.005 increase in entropy per decade of age (*p*-value < 2.2e-16), and a stark contrast with the diverging sites, which significantly *decrease* entropy with age, and could be considered “anti-entropic” since they become *more* predictable with age (**Figure 5B**). We validated these results in a second dataset, GSE128235, and found highly concordant results in both the proportion of DMPs that converge to (72%) and diverge from (28%) the mean with age, but also the significant increase in entropy at the converging sites of 0.005 per decade of age (*p*-value < 2.2e-16), and significant decrease in entropy of -0.004 per decade of age at the diverging sites (*p*-value < 2.2e-16).

**Figure 5.**
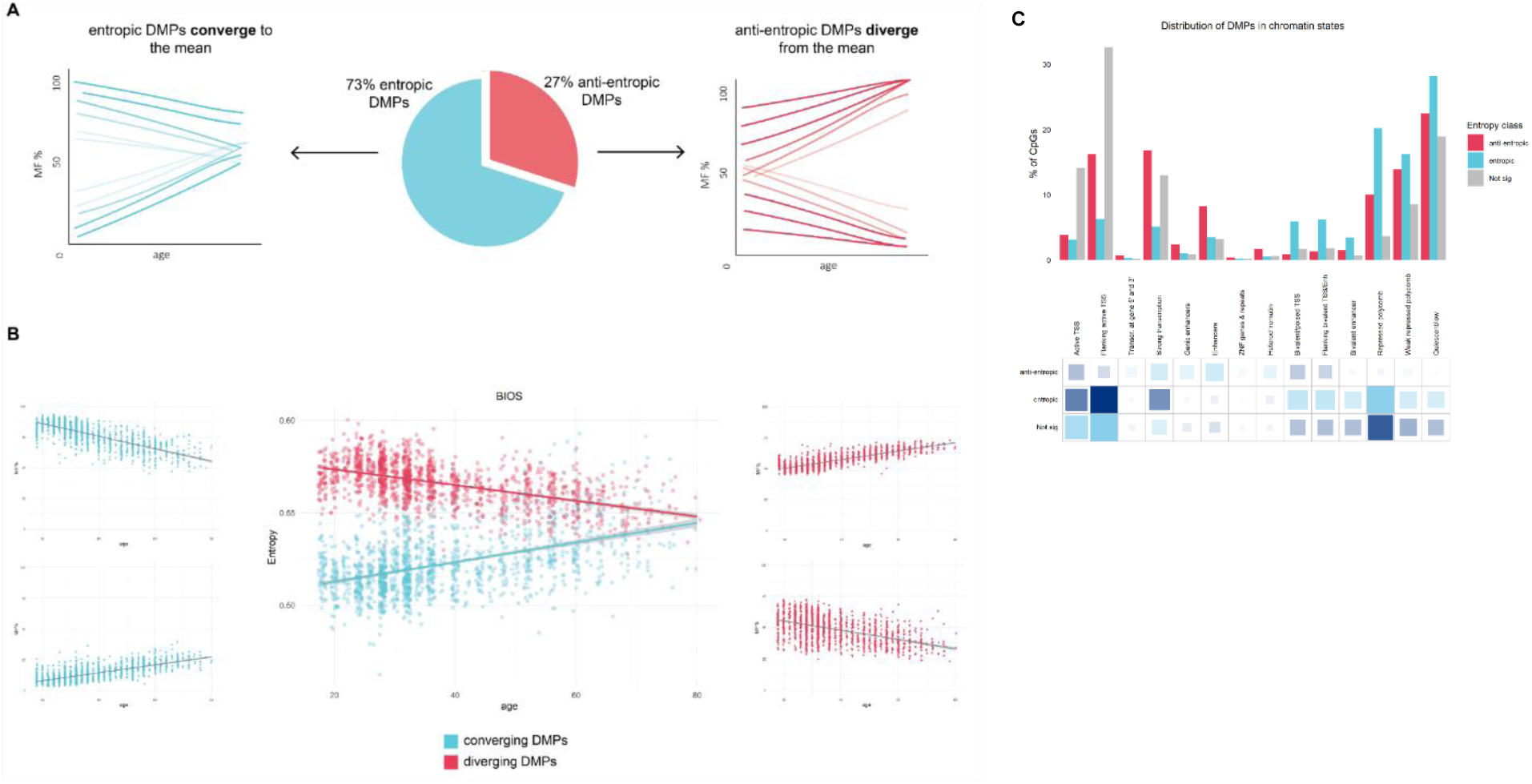
Contribution of differential methylation to changes in entropy with age. A) A pie chart of the proportion of DMPs that are entropic and converge to the mean (blue) and the proportion of DMPs that are anti-entropic and diverge from the mean (red) for the BIOS dataset in blood. The arrows point to hypothetical graphs that illustrate the aggregate regression lines for the CpGs that converge (left) and diverge (right) with age. B) In the far left (blue) and far right (red) panels are the methylation plots for two CpG sites, respectively, that are highly or lowly methylated in young (blue) and converge to the mean with age, and two methylation plots that are intermediately methylated in young (red) and diverge from the mean with age. C) Enrichment of entropic vs anti-entropic DMPs in chromatin states (A), biological pathways (B) and epigenetic clocks (C). A) Chromatin state enrichment of entropic and antientropic DMPs. A) Distribution of entropic (blue) and anti-entropic differentially methylated positions (DMPs) (red) and non-DMPs (grey) in chromatin states from peripheral blood mononuclear cells (PBMCs); The grids under the graph represent the residuals from the ꭓ2 test, with the size of the blocks in the grid being proportional to the cell’s contribution. Sky blue indicates over-representation or enrichment in the chromatin state, and navy represents under-representation or depletion in the chromatin state.

We then investigated the distribution of the entropic DMPs, anti-entropic DMPs and non-DMPs in chromatin states of PBMCs (**Figure 5A**), noting that entropic DMPs are overrepresented in bivalent promoters and enhancers, regions bound by Polycomb proteins, and quiescent states (ꭓ^2^ test *p-*value < 2.2e-16). In contrast, anti-entropic DMPs are overwhelmingly found at regions of strong transcription and enhancers (ꭓ^2^ test *p-*value < 2.2e-16) (**Figure 5B**).

We hypothesised that cell type heterogeneity would bias age-related changes in entropy estimates *upwards* (i.e. the increase in entropy with age would be inflated because of changes in cell type % with age). We first repeated the analyses after adjusting the DNAm profiles for blood cell types in each dataset (see Methods). There was a moderate correlation of 0.53 (*p*-value = 2.9e-5) between the effect sizes (i.e. the change in entropy per decade of age) before vs after adjustment for cell type proportions (**Extended Data Fig. 8A**), and half of the datasets displaying effect sizes that declined in magnitude after adjusting for cell type proportions (**Extended Data Fig. 8B**). However, the overall meta-analysis effect size remained unchanged after adjustment (0.0005 change in entropy per decade of age) (**Extended Data Fig. 8C**).

We then looked at age-related changes in entropy in datasets containing isolated cell types, speculating that if age-related changes in entropy were solely driven by changes in cell type %, we would fail to see an increase in entropy in these datasets. We looked at monocytes (GSE56046), CD4+ T cells (GSE59065, GSE56581 & GSE137593), CD8+ T cells (GSE59065), and B cells (GSE137594) (). In nearly all datasets, we failed to detect any change in entropy during ageing, but both CD4+ T and CD8+ T cells in GSE59065 displayed a marked increase in entropy during ageing (). Finally, we took advantage of the unique design of dataset GSE184269 that contains both mixed and sorted blood cells from the same individuals (naïve B cells, naïve CD4+ T cells, naïve CD8+ T cells, NK cells, monocytes and granulocytes in GSE184269), speculating that if age-related changes in entropy were solely driven by changes in cell type %, PBMCs would show higher entropy levels than sorted cell types. Entropy was markedly higher in PBMCs than NK, naïve CD4+ T, naïve CD8+ T and naïve B cells, but the highest entropy levels were in the heterogeneous class of granulocytes that comprise basophils, eosinophils, and neutrophils (**Extended Data Fig. 9**). These results suggest that changes in cell type composition during ageing partially account for age-related increases in entropy.

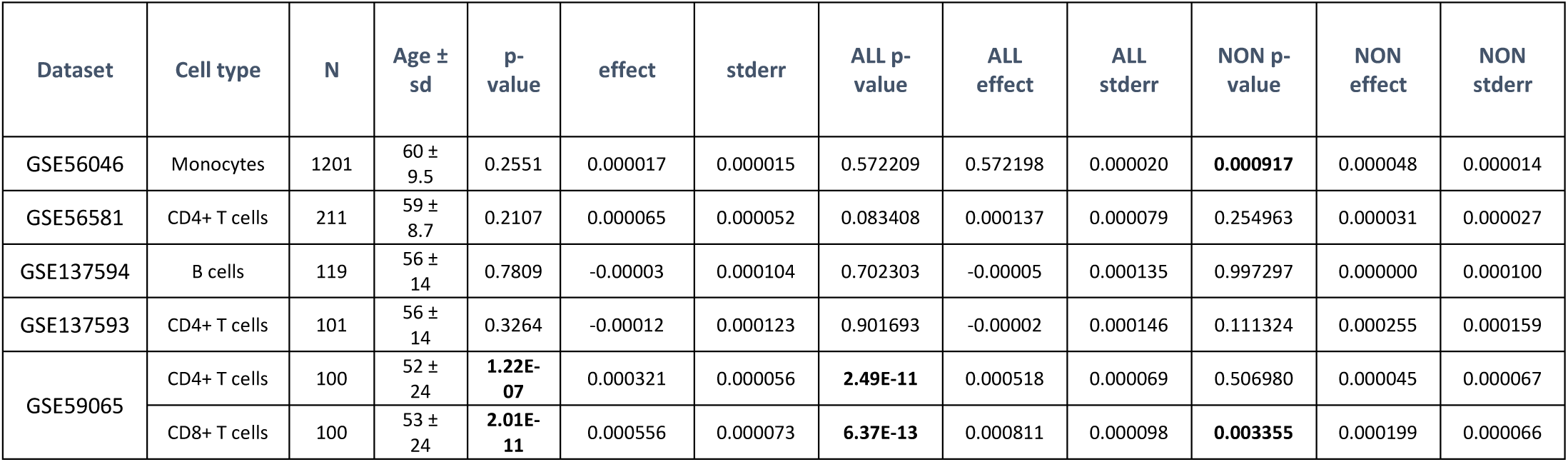

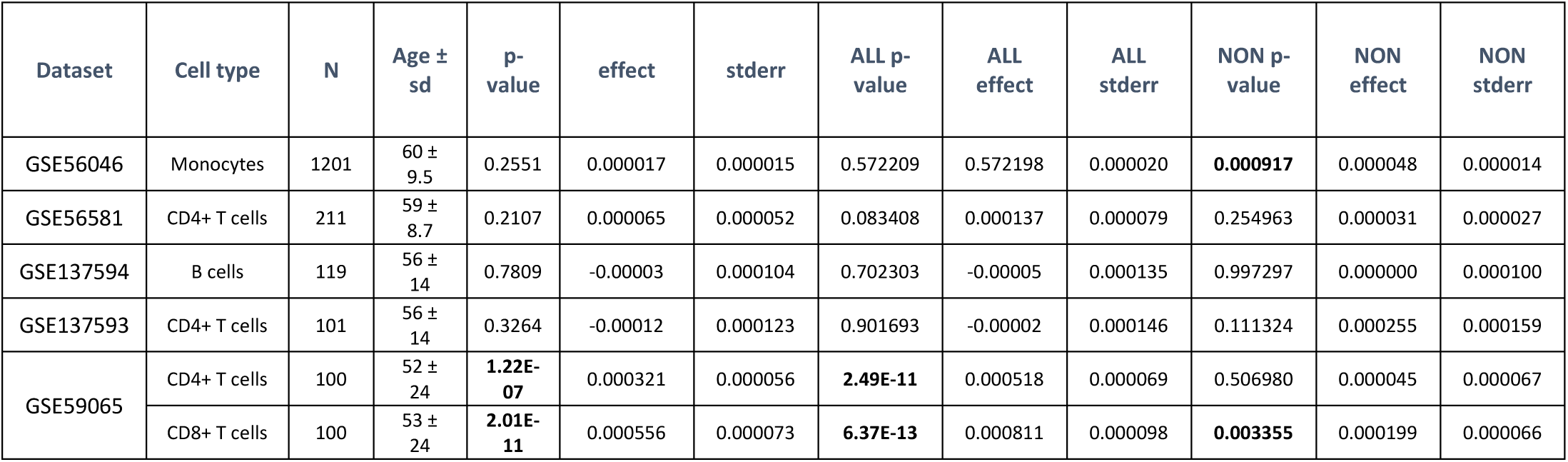

## Discussion

We demonstrated that during ageing, large proportions of the blood methylome slowly shift in mean methylation levels, and about half of them also become increasingly divergent between individuals as they get older. Our results are robust and replicable, owing to the unparalleled statistical power generated from > 32,000 samples in 56 independent cohorts. Importantly, these age-related changes occur largely independently of changes in the relative proportions of different blood cell types during ageing. Considering the widespread distribution of these DNAm changes, ageing appears to affect most molecular pathways, from development, metabolism, and cell-signalling to homeostatic processes. Shannon entropy increases in the blood methylome during ageing due to the presence of DMPs, and because the majority of DMPs trend towards intermediate methylation levels of 50%. This implies that during ageing, it becomes increasingly difficult to predict the methylation state of any given blood cell, in agreement with the smoothening of the epigenetic landscape. However, the overall increase in entropy is small in magnitude, and we uncovered that roughly one third of DMPs, particularly those enriched at enhancers and regions of active transcription, are ‘anti-entropic.’

While we know from previous studies that DMPs are a feature of ageing in blood^4,24–29^, our findings are unparalleled, revealing the sheer magnitude and omnipresence of these differential shifts that may have been previously underappreciated. Moreover, our study is 38 times larger than the largest known study of VMPs in blood^7^. Unlike previously reported^7^, we noted a significant overlap between DMPs and VMPs (i.e. many CpGs change in both average methylation and variance with age). We also show that the increase in entropy is driven by the differential shifts that trend from high and low methylation fractions in young, to intermediate methylation states at older ages; also referred to as the ‘smoothening of the epigenetic landscape,’ however, intermediately methylated CpGs that drift *away* from the mean towards fully methylated and unmethylated states decrease entropy, exhibiting ‘antientropic’ properties. It has been proposed that ‘anti-entropic’ CpGs could represent genes that become increasingly regulated^6^, a theory worth investigating further. Since the majority of DMPs are entropic, the net global effect is an increase in entropy with age.

In alignment with previous studies, we identified key processes related to development, differentiation and cell-signalling are altered during ageing^3^. The promoters and bodies of actively transcribed genes were fairly unaffected by epigenetic aging. In contrast, our results support previous evidence of age-related gains in methylation *and* variance at important regulatory regions, particularly bivalent domains that harbour both active and inactive histone marks and regions repressed by the Polycomb complex^3^. Bivalent regions prepare key developmental genes to be switched on during differentiation in specific cells, such as embryonic stem cells^30^. The epigenetic clock starts ticking upon differentiation^31^, and some gains in methylation may lead to loss of haematopoietic stem cell plasticity. However, these age-related epigenetic changes could affect all cell types in a tissue, not just stem cells. Bivalent promoters and poised enhancers are not exclusive to stem cells, and bivalency has been proposed to be a major reversible signature that distinguishes between cell types in adult tissues^30^. This would make sense, as epigenetic clocks work well in a plethora of tissues, even those that do not have many known stem cells^32^. Age-related hypermethylation may therefore hinder *all* cells from robustly maintaining their identity into old age; a process *shared* among individuals during chronological ageing, but that happens at different rates between people. Age-related loss of cellular identity has been previously reported in gene expression signatures in different somatic tissues of mice, whereby the transcription profiles *diverge* during development and *converge* during ageing^33^. A similar idea is described in a recent study as “ex-differentiation,” whereby aged cells lose their ability to maintain their identity due to epigenetic changes at developmental genes^34^. It is therefore plausible that environmental insults (i.e. sedentary behaviour, sun exposure, toxins, inflammation, injury) could accelerate or exacerbate these changes via metabolic stress, damage to the ECM and mitochondrial biogenesis^35^, and these would be detected as VMPs. The stochasticity or variability may be a good indicator of cumulative damage of the environment or non-specific damage, whereby the epigenome is adapting to changing environmental cues^36^. It is therefore somewhat puzzling that both chronological and biological clocks were strongly enriched for those CpGs that show changes in average *and* variance during aging (DMPs-VMPs). Intuitively, chronological clocks, whose objective is to predict time elapsed since birth, should contain an overwhelming number of homoscedastic DMPs as those carry less “noise” between individuals as they get older. Furthermore, the age-related sites we identified were not enriched in pace-of-aging clocks: while those clocks do not measure biological age per se and rather the speed at which one ages across different organ systems, we expected some overlap with age-related sites as the biological processes controlling the rate of aging should be somewhat related to the biological processes controlling biological age.

In contrast to age-related hypermethylation that was accompanied with large divergence between individuals, age-related hypomethylation was more homogeneous across the lifespan and mainly affected quiescent regions, with unclear functional consequences. It may reflect tissue-specific operations^3^, and comparing our results across other tissues would be necessary to confirm this. In blood, hypomethylation has been hypothesised to be immunogenic and contribute to inflammaging^37^. Our pathway enrichment analysis revealed DMGs and VMGs to be associated with abnormal inflammatory cascades, and multiple organ system abnormalities, consistent with chronic inflammation as a hallmark of ageing^2^. This inflammation is systemic and associated with various ageing-related phenotypes in multiple tissues.

Cellular heterogeneity in tissues poses a significant challenge in DNAm studies^8^. We found a high overlap of significant CpGs in both the adjusted and unadjusted analyses of DMPs and to a lesser extent, VMPs. These represent age-associated DNAm changes that are either present in all underlying subtypes, or only in a predominant subtype^8^, a finding that overlaps with previous studies^7,38^. While majority of the age-associated changes seem independent of changes in cell composition, about one third of VMPs were significant *only* in the unadjusted analyses, likely capturing variability related to changes in cell composition with age. It has been suggested that the increase in ‘transcriptional noise’ underlying a loss of cell identity lacks sufficient evidence and may instead be related to the age-related changes in cell composition^37^. As such, these VMPs may be capturing an important phenotype of ageing in blood (and potentially other tissues), such as the remodeling of the immune system cell type composition^39^. Importantly, performing cell type correction is recommended *in addition* to an unadjusted analysis^8^, particularly in ageing studies, since increasing cellular heterogeneity could reflect an important biological process or age-related phenotype. Future work should also endeavor to identify age-related changes *within* individual cell types^40^, to understand the relative contribution of different cellular compartments in age-related diseases.

The significance of our cohort substructure, comprising extensive individual datasets with samples spanning a wide age range (6 - 101 years old), was evident when attempting to detect all kinds of age-related changes. Detecting VMPs using the Breusch-Pagan test for heteroscedasticity necessitated not only large sample sizes and a broad age range, but also possibly individuals at the ‘upper limit’ of old age. Interestingly, a recent preprint in mouse blood showed that the variance of DNAm increased only at very old ages^36^, implying that previous human studies may have been limited due to small sample sizes, narrow age ranges, and a scarcity of very old individuals, hindering investigations into variance or stochasticity with age^28^. In contrast, the standard linear model used to detect DMPs is less sensitive to the characteristics of the dataset substructure. However, the age range in the datasets was skewed, with varying sex distributions, diseases statuses, and other factors that could influence the effect of age on the methylome. For instance, age-related changes might be more pronounced in different ethnicities or sexes, introducing some variability. Given the substantial sample size, we see this heterogeneity as a strength more than a weakness, because it implies that the DMPs, VMPs and entropy we detected hold true for a broad range of individuals and could be considered “universal” markers of ageing. One caveat to note is that the ethnic representation of the samples was overwhelmingly white, clearly highlighting the need to obtain a better representation of ethnicities in DNAm profiles.

In conclusion, using an unparalleled cohort of methylomes in humans, we provide a comprehensive picture of the global age-related changes observed in blood. Future work could focus on assessing the effect of longevity interventions, such as exercise, nutrition, supplementation, etc., on these age-related signatures *in vivo* humans.

## Methods

This study was conducted as a large-scale, multi-tissue EWAS meta-analysis of age. Bioinformatics techniques were applied to analyse and interpret large amounts of existing epigenomic data. By exploiting the power of meta-analysis, we overcome many limitations of ‘omics’ research. Specifically, very large sample sizes are required to detect changes with small effect sizes, which is the case of age-related changes in DNAm profiles^41^. Our approach was therefore robust for identifying subtle, yet highly reproducible shifts in DNAm that accrue over chronological time in a wide variety of populations (e.g. males/females, conditioned/healthy individuals, etc.).

### Data mining

To create the repository of blood DNAm datasets, we have carried out a comprehensive data mining enterprise collecting the methylomes of 32,136 human samples from 56 datasets, profiled on the Illumina Methylation array platforms (27K, 450K and EPIC) (**Figure 1**, **Supplementary Table 1**). This includes 49 open-access datasets from Gene Expression Omnibus (GEO) repository and 1 from ArrayExpress, 4 datasets from the controlled access database of Genotypes and Phenotypes (dbGaP), 1 dataset from the controlled access European Genome-Phenome Archive (EGA), and 1 dataset from a controlled access independent repository (**Supplementary Table 1**). Datasets with fewer than 30 samples or an age standard deviation < 5 were excluded, as low age variability and low sample size severely impairs the ability of the linear models to detect age-related patterns reliably. Samples with a cancer diagnosis were also excluded, as cancer samples show highly unusual DNAm patterns that would likely skew the analysis^12^.

### Pre-processing

Datasets with raw DNAm data available were pre-processed, normalised, and filtered using the R statistical software (www.r-project.org), and following the *ChAMP*^11^ pipeline. Methylated and unmethylated signals or IDAT files were used for the pre-processing. In accordance with the default parameters of the *champ.load* function, any sample with >10% of probes with a detection *p-*value > 0.01 was removed^20^. All probes with missing β-values (a detection *p-*value > 0.01), with a bead count < 3 in at least 5% of samples, non-CG probes and probes aligning to multiple locations were filtered out. Probes located on the sex chromosomes were also filtered out in datasets containing both males and females, as well as probes mapping to single nucleotide polymorphisms (SNPs)^11^. Additional cross-hybridising probes identified by Pidsley *et al*.^42^ were also filtered out^7^. The methylation β-values obtained were calculated as the ratio of the methylated probe intensity and the overall intensity, as follows:

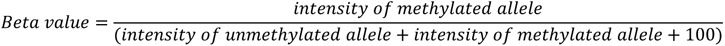

The Type I and Type II probe designs that are generated from the 450K and EPIC Illumina arrays were normalised using the *champ.norm* function^20^. We explored the technical and biological sources of variation in each dataset using a singular value decomposition method provided by the *champ.SVD* function^20^. The *ComBat* function from the *sva* package was used to adjust for technical variation from the slide and position on the slide if this information was available^43^. We could not perform batch correction if this information was unavailable. Finally, samples whose annotated sex was discordant with predicted sex (according to the *getSex* function from the *minfi* package), were removed^44^. Any missing information required for pre-processing, including raw IDAT files, batch information, detection *p*-values or the age of the samples, was requested from the corresponding authors at the time of pre-processing (**Supplementary Table 1**). We used the pre-processed matrices for datasets that we were unable to preprocess ourselves due to missing information (**Supplementary Table 1**).

### Statistical framework

#### DMPs

To identify DMPs, we performed an EWAS in each dataset using linear regression models and moderated Bayesian statistics, as implemented in the *limma* package^45,46^. A linear model was fitted for each dataset independently. DNAm was regressed against age and other dataset-specific covariates (i.e., sex, batch, body mass index (BMI)) (**Supplementary Table 1**), when this information was available (See below for the inclusion/exclusion criteria of covariates). If a dataset included repeated measures of the same individual or related individuals (e.g., twins), we added a random effect to the model (using the *duplicateCorrelation* function of the *limma* package). Linear models were performed using M-values (a logit transformation of β-values), which have more favourable statistical properties for differential analysis^47^.

To then compare the effect of cell type heterogeneity on age-associated DNAm changes, we repeated all linear models adjusting for cell type proportions in whole blood. We applied the *champ.refbase* method, as implemented in the *ChAMP* package, to estimate the cell type proportions for granulocytes, monocytes, NK cells, CD4+ T cells, CD8+ T cells and B cells, for each sample^20^. We then repeated the linear model for each blood dataset including the 5 largest cell types to remove the confounding effect of cell type proportion on DMPs.

To assess whether the direction of DNAm with age (hypo/hyper) depends on the baseline DNAm levels, we classified DMPs in “high”, “intermediate” and “low” methylation categories using a large dataset (BIOS). In this dataset and for each DMP, we calculated the average MF and classified a DMP as ‘high’ if MF >= 75%, ‘low’ if MF <= 25%, and the remainder ‘intermediate’. This categorisation was made separately for young (i.e. < 30 y.o) and old (> 60 y.o.) individuals, and we assessed whether the DMP trended from ‘high’ in young to ‘intermediate’ or ‘low’ in old, etc.

#### VMPs

Age-associated VMPs were identified using the Breusch-Pagan test for heteroscedasticity, which is a two-way regression that models the change in DNAm variance as a function of age (i.e. it tests if the variance in DNAm levels (adjusted for covariates) is dependent on age)^3,48^. DNAm was first regressed against age and other confounders for each dataset (i.e. the linear model to identify DMPs) to obtain residuals. The residuals were then extracted from this model and squared. We ran the Shapiro-Wilk test to remove CpGs where residuals strongly deviated from normality (i.e. DNAm sites that are associated with SNPs not filtered out during pre-processing)^4^, and subsequently regressed the squared residuals of the remaining markers against age. All analyses were repeated and corrected for cell types to compare the effect of cell type heterogeneity on age-associated VMPs.

#### Entropy

Shannon entropy ranges between 0 and 1, taking its maximum value when the methylation fraction in a given set of CpGs, measured over a population of cells, is 50%^49^. Shannon entropy was calculated for each sample in each dataset, using a probability formula adapted to handle DNAm data^3,4,7^.

To calculate the genome-wide Shannon entropy, a linear model is fitted for each dataset on the logit transformed M-values, adjusting for dataset covariates where appropriate (e.g. sex, BMI, batch). Age was not included in this model, as we did not want to remove the effect of age on the β-values. The *mean* M-values from the original matrix were added to the residuals from the linear model, and then transformed back to obtain *adjusted* β-values, as the Shannon entropy formula has been adapted specifically to handle β-values.

Using the adjusted β-values, genome-wide Shannon entropy was computed for each sample in each dataset according to the formula^3,4^:

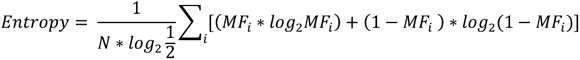

where *MF_i_* is the methylation fraction (e.g. beta value) for the *i^th^* CpG probe and *N* is the total number of CpGs.

To calculate the effect of age on Shannon entropy, a linear model was fitted for each dataset regressing age against entropy and recorded the summary statistics for each dataset (**Supplementary Table 2**), as follows:

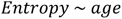

Since genome-wide entropy captures the complexity of the entire system in a single measure, we sought to determine the contribution of the various features of ageing that may be driving genome-wide changes in entropy. To do so, we then repeated the analysis by calculating entropy for each sample, in each dataset, in each tissue, on all the *age-associated CpGs* identified from the meta-analysis in aims 1 and 2 (i.e. a complete list of both DMPs and VMPs), as well as the *non-age-associated CpGs* (i.e. the complete list of non-DMPs and non-VMPs). We then repeated the above steps calculating entropy CpGs that were only *DMPs*, CpGs that were both *DMPs and VMPs*, CpGs that were only *VMPs* (**Supplementary Table 3**), *entropic DMPs*, and *anti-entropic DMPs*. We also re-calculated entropy on all measurements and corrected for cell type heterogeneity, to compare the effect of cell type composition on age-associated changes in entropy (**Supplementary Table 2**), as follows:

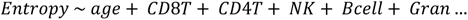

In addition, we calculated entropy for datasets from sorted cell types on the genome-wide set of CpGs, only on the age-associated CpGs, and on the non-age-associated CpGs (**Supplementary Table 4**).

#### Adjusting for covariates

We took careful consideration when adjusting for disease-specific covariates in our linear models to ensure that we did not unnecessarily remove the effect of age on DNAm.

While no official classification of ‘age-related’ exists, 92 conditions have been classified as ‘age-related’ based upon an *exponential* increase in incidence with age using a two-step mathematical modelling technique^50^. To have a direct view of the relationship between age and each disease/condition present in our datasets (Supplementary Tables 1 - 6), we used the GDB data exploration tool (https://vizhub.healthdata.org/gbd-compare/).

Based on their classification and the GDB tool, we *included* the following conditions/phenotypes as covariates that show no relationship with age, including depression, anaemia, inflammatory bowel disease / Crohn’s diseases, lupus, non-alcohol steatohepatitis (NASH) / non-alcoholic fatty liver disease (NAFLD) and asthma, and *excluded* conditions/phenotypes that show a relationship with age (either increase or decrease in prevalence with age), including progressive supranuclear palsy, Alzheimer’s disease/dementia, cardiovascular disease, ischemic stroke, Parkinson’s disease, COVID-19, cardiomyopathy, chronic obstructive pulmonary disease (COPD), schizophrenia, anxiety disorders, osteoarthritis, rheumatoid arthritis, multiple sclerosis, psoriasis, type 2 diabetes and cirrhosis.

Importantly, some diseases did show a relationship with age, but cannot be considered age-related as they don’t depend on age-related functional decline of the body, but on age-related differences in behaviour (e.g. HIV infection, drug use disorder), and were included as covariates. In addition, factors that are not age-related but accelerate or slow down ageing (e.g. smoking, BMI, hypertension, exercise training and bariatric surgery) were also included as covariates.

#### Meta-analyses of DMPs and age

As described above, an EWAS of age was performed in each dataset independently. Each EWAS was adjusted for bias and inflation using the empirical null distribution as implemented in *bacon*^41^. The results from the independent EWAS were then pooled, using an inverse variance based fixed-effects meta-analysis implemented in METAL^51^. This approach computes a weighted average of the results, using the individual effect size estimates and standard errors extracted from each independent EWAS. The overlap in CpGs between datasets was imperfect (not all CpGs were present in all datasets), as we used three different Illumina array platforms, and since different CpGs are filtered out during the pre-processing of individual datasets. We restricted our analysis to CpGs that were present in at least three datasets. Age-associated DMPs were then identified using a stringent meta-analysis false discovery rate (FDR) < 0.005. The meta-analysis was repeated with datasets adjusted for cell types.

#### Meta-analyses of VMPs and age

As for the DMPs, we used METAL to pool results from the independent EWAS, but we followed a *sample size*-based fixed effects meta-analysis (instead of an inverse-variance method)^51^ . This approach was more appropriate to meta-analyse the χ^2^ test statistic that is the output of the Breusch-Pagan test; it relies on the *sample size* of each dataset and the *p*-value at each CpG. We restricted our analysis to CpGs that were present in at least 15% of the samples. Age-associated VMPs were identified using a stringent meta-analysis FDR <0.005. The meta-analysis was repeated with datasets adjusted for cell types.

#### Meta-analyses of entropy and age

To identify the change in entropy with age in each tissue, the summary statistics (i.e. effect size and standard error) extracted from the independent entropy regressions were pooled using a fixed effects meta-analysis using the R package *metafor*^52^ (**Supplementary Table 2**). We meta-analysed the summary statistics for the genome-wide entropy results, as well as for the age-related CpGs, and the non-age-related CpGs. All meta-analyses in blood were repeated for the cell-type corrected analyses, to compare the effect of cell heterogeneity on entropy.

#### Chromatin state enrichment, pathway analysis and epigenetic clock CpGs analysis

To aid in the biological interpretation of the identified age-associated methylation sites, we tested whether DMPs and VMPs showed any enrichment in chromatin states. This was done by comparing the distribution of the blood-specific VMPs and DMPs with that of non-VMPs and non-DMPs, respectively, in the different chromatin states profiled in peripheral blood mononuclear cells (PBMCs) from the Roadmap Epigenomics Project with a Fischer’s exact test^10^.

To gain insights into the cellular and phenotype consequences of ageing on the blood methylome, we tested whether genes belonging to gene ontology (GO) terms (GO gene set in MsigDB), human phenotype ontologies (HPO gene set in MsigDB), canonical pathways (CP gene set in MsigDB), expression signatures of genetic and chemical perturbations (CGP gene set in MsigDB) and immunological signatures (C7 gene set in MsigDB) were enriched among the VMPs and DMPs using the *gsameth* function from the *missMethyl* package^53^. An improved adaptation of Zhou et al.’s comprehensive annotation was used to assign one or more genes to each VMP and DMP^11^ All GO, HPO, CP and CGO terms were deemed significant at an FDR < 0.005^54,55^.

We also tested whether DMPs and VMPs were particularly over- or under-represented in CpGs that make up the different epigenetic clocks that were developed using elastic net. We tested clocks trained to predict chronological age (Horvath’s pan-tissue clock, 353 CpGs^12^; Hannum’s clock, 71 CpGs^4^; the blood clock developed by Zhang et al. 2019, 514 CpGs^13^; the centenarian clock, 747 CpGs^14^; version 2 of the mammalian universal clock, 816 CpGs ^15^); biological age (PhenoAge, 513 CpGs^16^), and the pace of aging (DunedinPoAm, 46 CpGs^17^; DunedinPACE, 173 CpGs^18^). This was done by comparing the distribution of the age-related CpGs with that of non-age-related CpGs, respectively, among the list of CpGs that make up different clocks, with the help of a χ^2^ test. We used a different background of probes for each clock, as each clock was developed from a different pool of CpGs (e.g. the pan-tissue clock was developed from a pool of ∼21,000 CpGs common to all Illumina HumanMethylation arrays, while DunedinPACE was developed from probes common to the 450K and EPIC arrays that showed good test-retest reliability). Note that we could not perform this enrichment for clocks whose list of CpGs is not open-access, such as GrimAge^56^, and for clocks that use *all* probes on the Illumina HumanMethylation arrays, such as Altum Age^57^, and PC-based clocks^58^. All χ^2^ test p-values were adjusted for multiple testing and only those tests with FDR < 0.005 were deemed significant.

#### Code availability

Code will be made available at the time of publication.

#### Figures

R studio ggplot2 from the tidyverse package and Cytoscape.

#### Data availability

Four datasets are sourced from the database of Genotypes and Phenotypes (dbGaP) and include The Genetics of Lipid-Lowering Drugs and Diet Network Lipidomics Study (GOLDN; accession number <u>phs000741.v2.p1</u>), the Women’s Health Initiative (WHI; accession number <u>phs001335.v2.p3</u>), the Normative Aging Study (NAS, accession number <u>phs000853.v2.p2</u>) and the Framingham Heart Study (FHS, accession number <u>phs000974.v5.p4</u>). The Biobank-based Integrative Omics Study (BIOS) can be requested and downloaded from the European Genome-Phenome Archive (EGA), accession <u>EGAS00001001077</u>. The Jackson Heart Study (JHS) can be requested from the <u>Jackson Heart Study website</u>. The Swedish Adoption/Twin Study of Aging (SATSA) is available from ArrayExpress under the accession number <u>E-MTAB-7309</u>. The remaining datasets are sourced from the Gene Expression Omnibus platform, including <u>GSE55763</u>, <u>GSE128235</u>, <u>GSE99624</u>, <u>GSE115278</u>, <u>GSE87571</u>, <u>GSE53740</u>, <u>GSE58045</u>, <u>GSE77445</u>, <u>GSE49904</u>, <u>GSE80417</u>, <u>GSE42861</u>, <u>GSE51032</u>, <u>GSE67705</u>, <u>GSE32148</u>, <u>GSE106648</u>, <u>GSE69138</u>, <u>GSE50660</u>, <u>GSE40279</u>, <u>GSE41037</u>, <u>GSE41169</u>, <u>GSE53840</u>, <u>GSE67751</u>, <u>GSE72775</u>, <u>GSE111629</u>, <u>GSE72774</u>, <u>GSE72776</u>, <u>GSE166611</u>, GSE164056.

## Acknowledgements

This work was supported by Nir Eynon’s National Health & Medical Research Council (NHMRC) Investigator Grant (APP1194159), as well as Sarah Voisin’s NHMRC Early Career Fellowship (APP1157732). The Gene SMART study is also supported by an Australian Research Council (ARC) Discovery Project Grants (DP190103081 & DP200101830 ).

The Jackson Heart Study (JHS) is supported and conducted in collaboration with Jackson State University (HHSN268201800013I), Tougaloo College (HHSN268201800014I), the Mississippi State Department of Health (HHSN268201800015I) and the University of Mississippi Medical Center (HHSN268201800010I, HHSN268201800011I and HHSN268201800012I) contracts from the National Heart, Lung, and Blood Institute (NHLBI) and the National Institute on Minority Health and Health Disparities (NIMHD). The authors also wish to thank the staffs and participants of the JHS.

<u>JHS disclaimer-</u> The views expressed in this manuscript are those of the authors and do not necessarily represent the views of the National Heart, Lung, and Blood Institute; the National Institutes of Health; or the U.S. Department of Health and Human Services.

This study makes use of data generated by the Biobank-based Integrative Omics Study. A full list of the investigators is available from http://www.bbmri.nl/en-gb/activities/rainbow-projects/bios. Funding for the project was provided by the Netherlands Organization for Scientific Research under award number 184021007, dated July 9, 2009 and made available as a Rainbow Project of the Biobanking and Biomolecular Research Infrastructure Netherlands (BBMRI-NL).

## Disclaimer

The views expressed in this manuscript are those of the authors and do not necessarily represent the views of the National Heart, Lung, and Blood Institute; the National Institutes of Health; or the U.S. Department of Health and Human Services.

---

**Extended Data Fig. 1.**
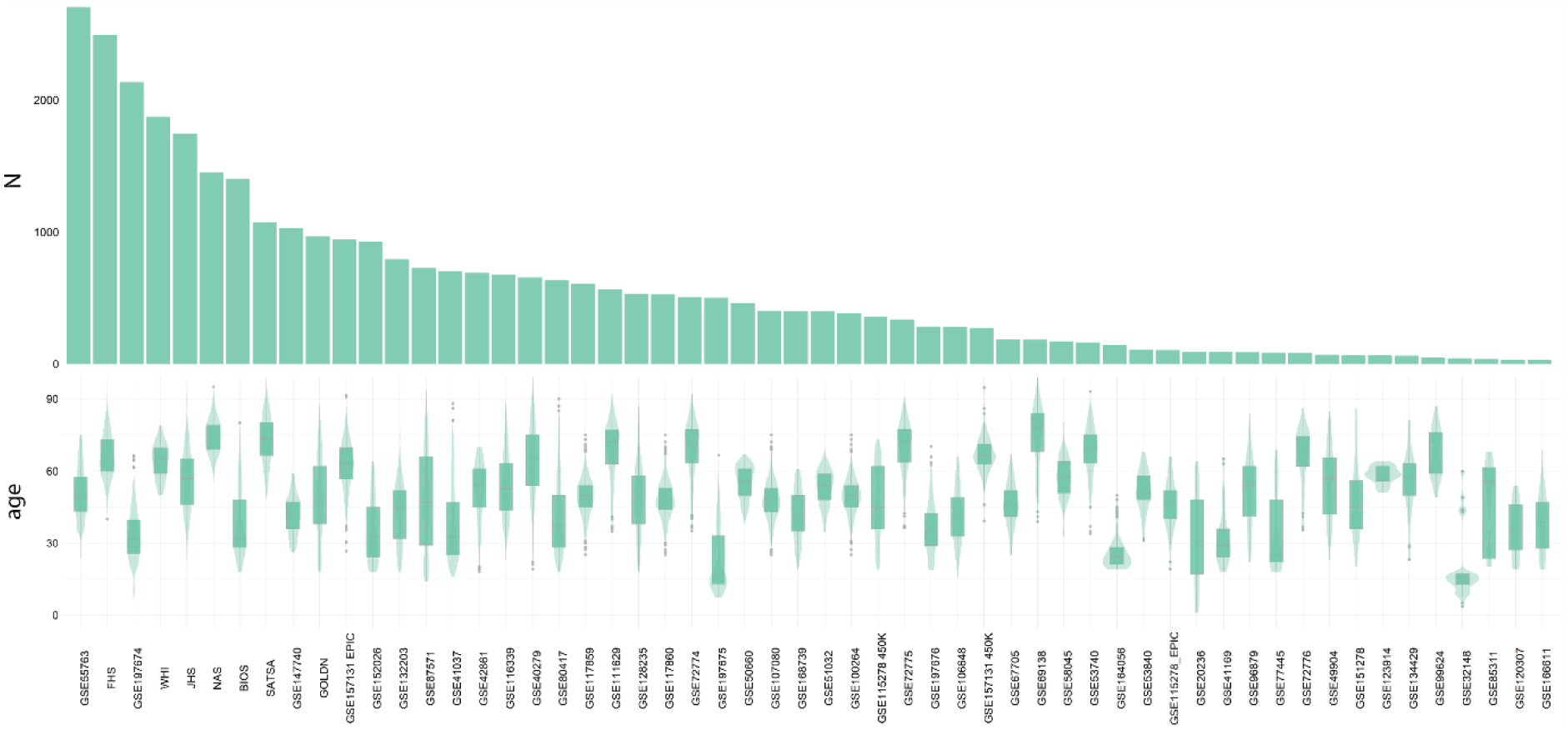
Sample size and age distribution in the 56 blood datasets analysed in this study. Bottom panel: boxplots & violin plots displaying the distribution of samples across the age range. Top panel: number of samples in each dataset (N), ordered from largest (left) to smallest (right).

**Extended Data Fig. 2.**
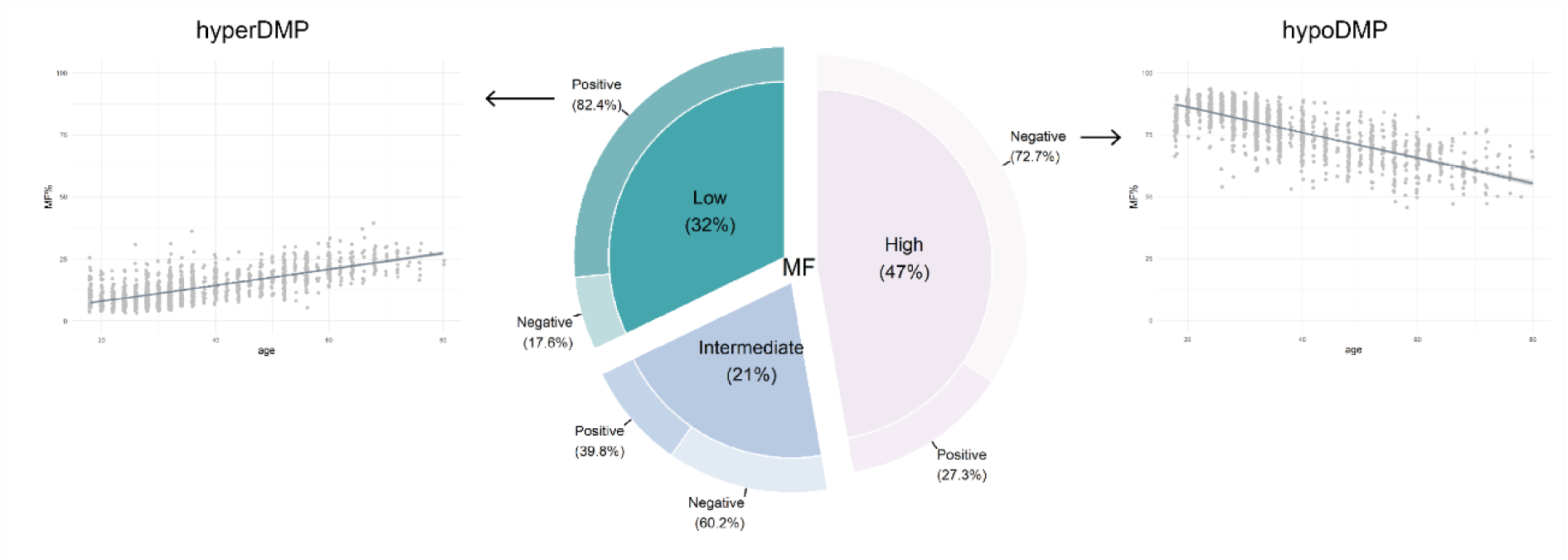
Doughnut chart of the proportions of highly, intermediately, and lowly methylated with age. The inner pie of the donut represents the proportions of CpGs that are highly methylated (<75%), intermediately methylated (25 – 75%) and lowly methylated (<25%) at baseline in a single blood dataset (BIOS), with the direction of change in methylation with age in the outer circle. A positive direction implies that a DMP increases in methylation with age, and a negative direction implies a DMP loses methylation with age. MF = Methylation Fraction

**Extended Data Fig. 3.**
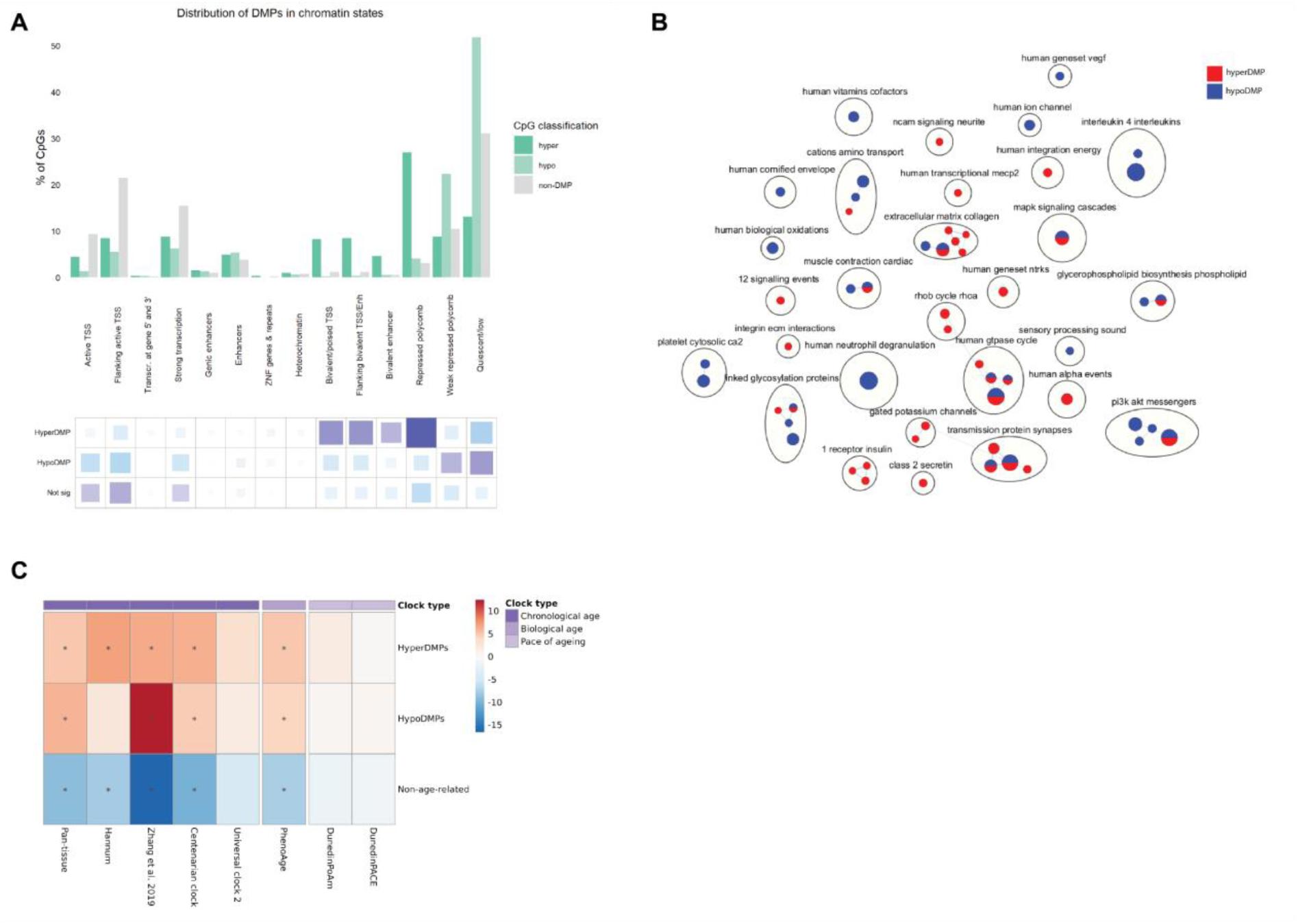
Enrichment of hypomethylated and hypermethylated differentially methylated positions (DMPs) and non-DMPs in chromatin states (A), biological pathways (B) and epigenetic clocks (C). **A)** Distribution of hypoDMPs, hyperDMPs and non-DMPs in chromatin states from peripheral blood mononuclear cells; the grids under the graph represent the residuals from the ꭓ^2^ test, with the size of the blocks in the grid being proportional to the cell’s contribution. Purple indicates over-representation or enrichment in the chromatin state, and blue represents under-representation or depletion in the chromatin state. **B)** Enrichment map showing Reactome pathways enriched in hypomethylated or hypermethylated DMPs. Nodes in the network represent pathways and similar pathways with many common genes are connected. Groups of similar pathways are indicated. Nodes are coloured according to significance in different DMPs types: blue (hypomethylated) and red (hypermethylated). **C)** Relative over-(red) or under- (blue) representation of clock CpGs in hyperDMPs, hypoDMPs and non-age-related CpGs.

**Extended Data Fig. 4.**
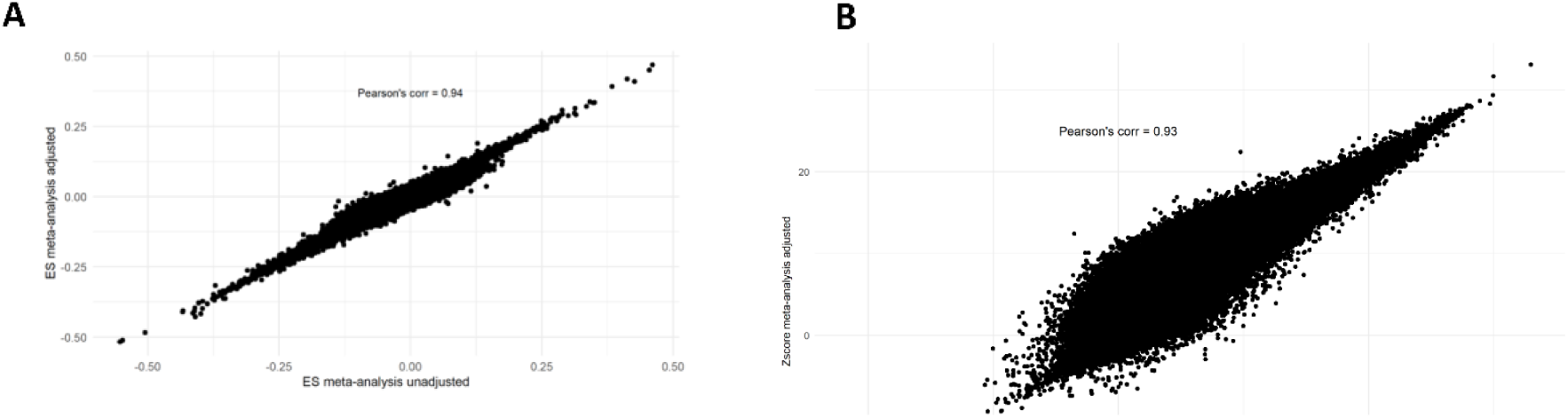
Comparison of DMP and VMP meta-analyses with and without correction for cell types. **A)** A correlation plot of the effect sizes of the CpGs meta-analysed in the cell type adjusted versus the meta-analysis not adjusted for cell types of differential methylation and age in blood. The effect sizes (ES) from the meta-analysis not adjusted for cell types on the x-axis and the ES for the meta-analysis adjusted for cell types on the y-axis. **B)** A correlation plot of the Zscores of the CpGs in the variable methylation and age meta-analyses in both the cell type adjusted and the unadjusted meta-analysis in blood. The Zscore from the meta-analysis not unadjusted for cell types on the x-axis and the Zscore for the meta-analysis adjusted for cell types on the y-axis.

**Extended Data Fig. 5.**
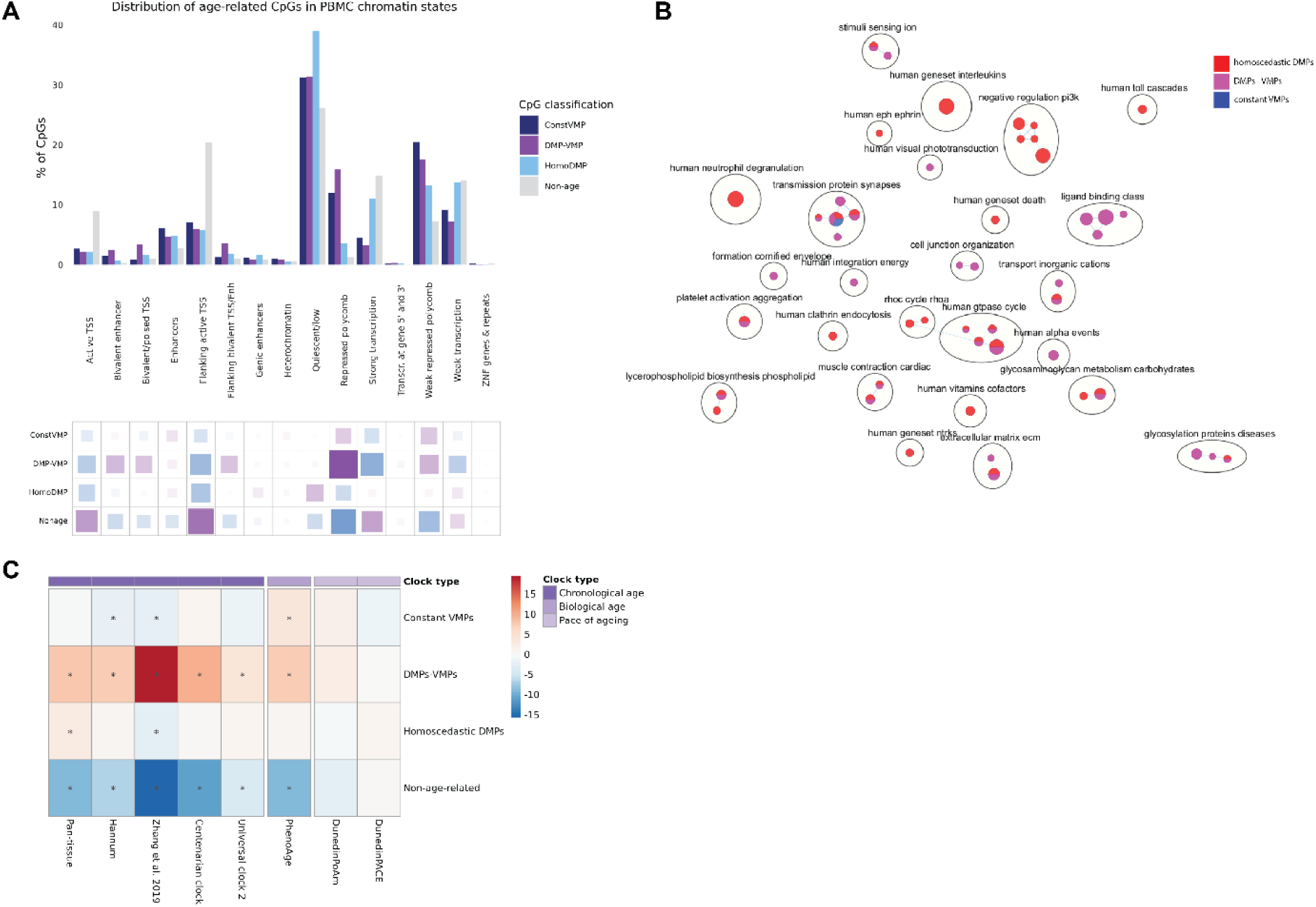
Enrichment of three classes of age-associated CpGs in chromatin states (A), biological pathways (B) and epigenetic clocks (C). **A)** Distribution of homoscedastic DMPs, DMPs-VMPs, constant VMPs and non-age-related CpGs in chromatin states from PBMCs. The grids under the graph represent the residuals from the ꭓ^2^ test, with the size of the blocks in the grid being proportional to the cell’s contribution. Purple indicates over-representation or enrichment in the chromatin state, and blue represents under-representation or depletion in the chromatin state. **B)** Enrichment map showing Reactome pathways enriched in homoscedastic DMPs, DMPs-VMPs, or constant VMPs. Nodes in the network represent pathways and similar pathways with many common genes are connected. Groups of similar pathways are indicated. Nodes are coloured according to significance in different age-related CpG types: red (homoscedastic DMPs), purple (DMPs-VMPs) and blue (constant VMPs). **C)** Relative over- (red) or under- (blue) representation of clock CpGs in homoscedastic DMPs, DMPs-VMPs, constant VMPs and non-age-related CpGs.

**Extended Data Fig. 6.**
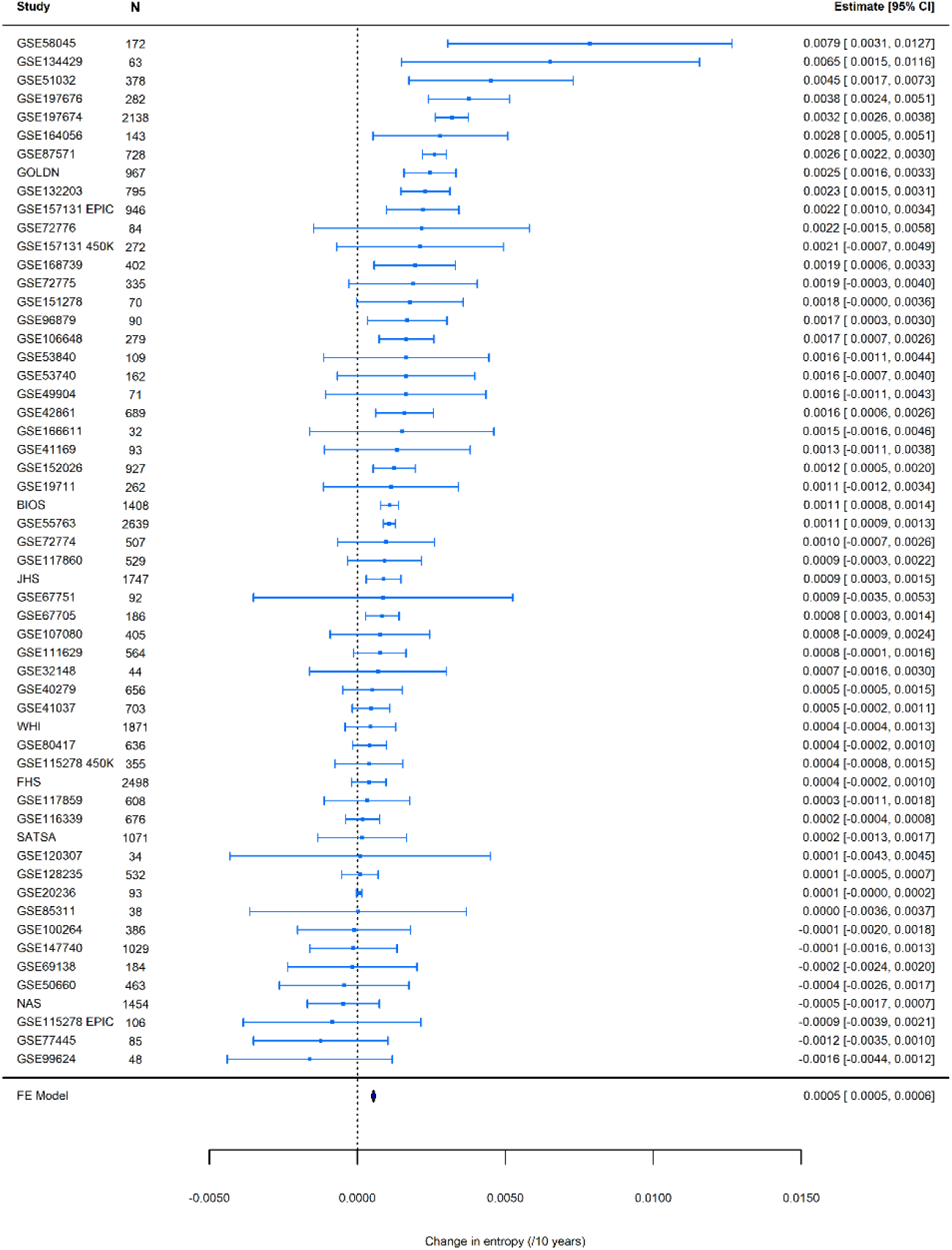
Forest plot of the genome-wide meta-analysis of entropy and age in blood. On the x axis is the change in entropy per decade of age, with the effect size and standard errors from each independent EWAS plotted on the y-axis. The dataset, sample size (N) and mean age ± SD (standard deviation) are on the left side of the plot and the 95% CI intervals are displayed on the right panel. The meta-analysis effect size is represented by the blue polygon.

**Extended Data Fig. 7.**
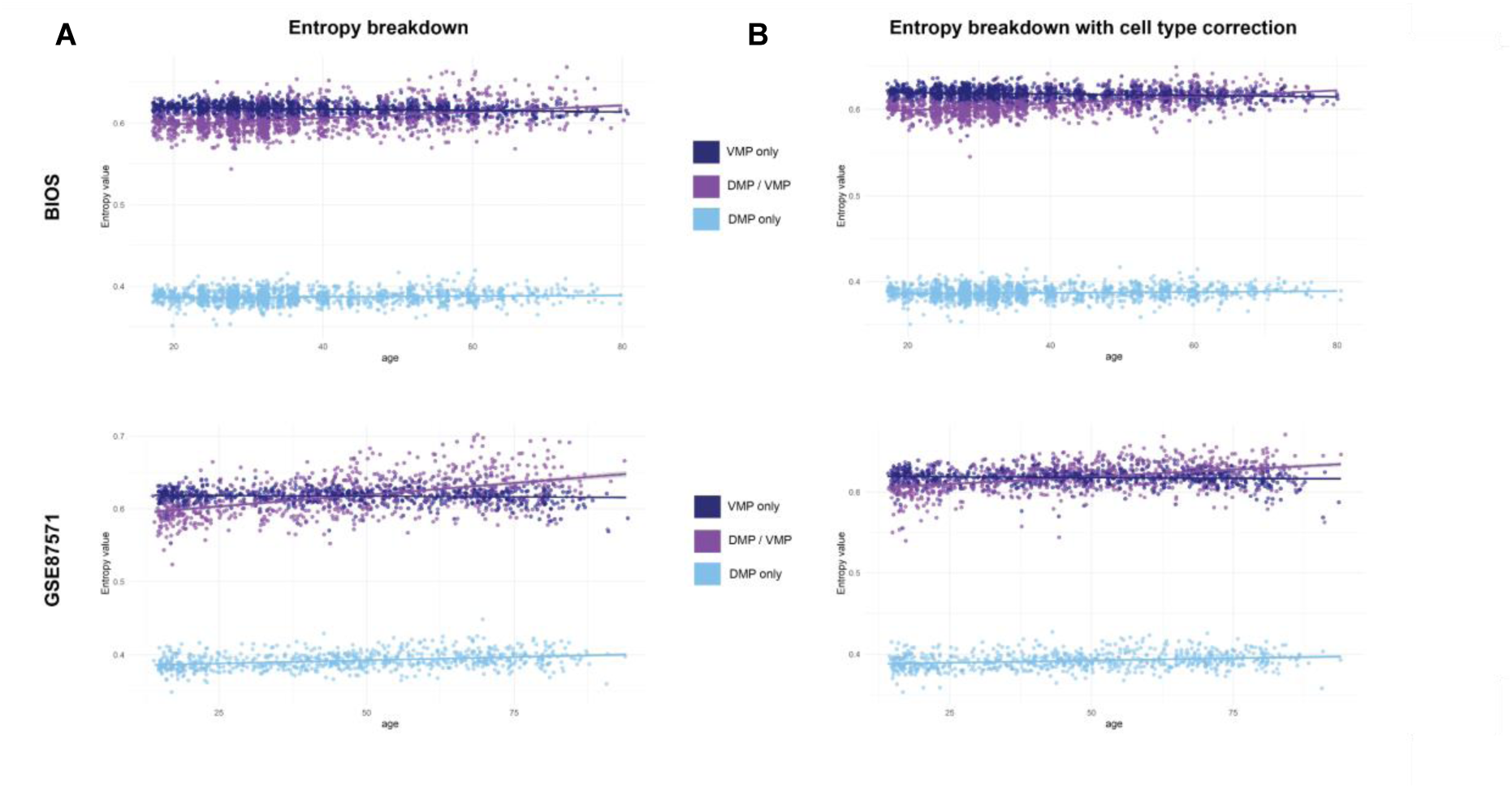
A comparison of the changes in entropy with age in homoscedastic DMPs, DMPs-VMPs and constant VMPs. Entropy was calculated for each type of age-related CpG on the y-axis and plotted against age (x-axis) for two datasets: BIOS and GSE87571. Analyses were performed A) unadjusted for cell types and B) adjusting for cell types.

**Extended Data Fig. 8.**
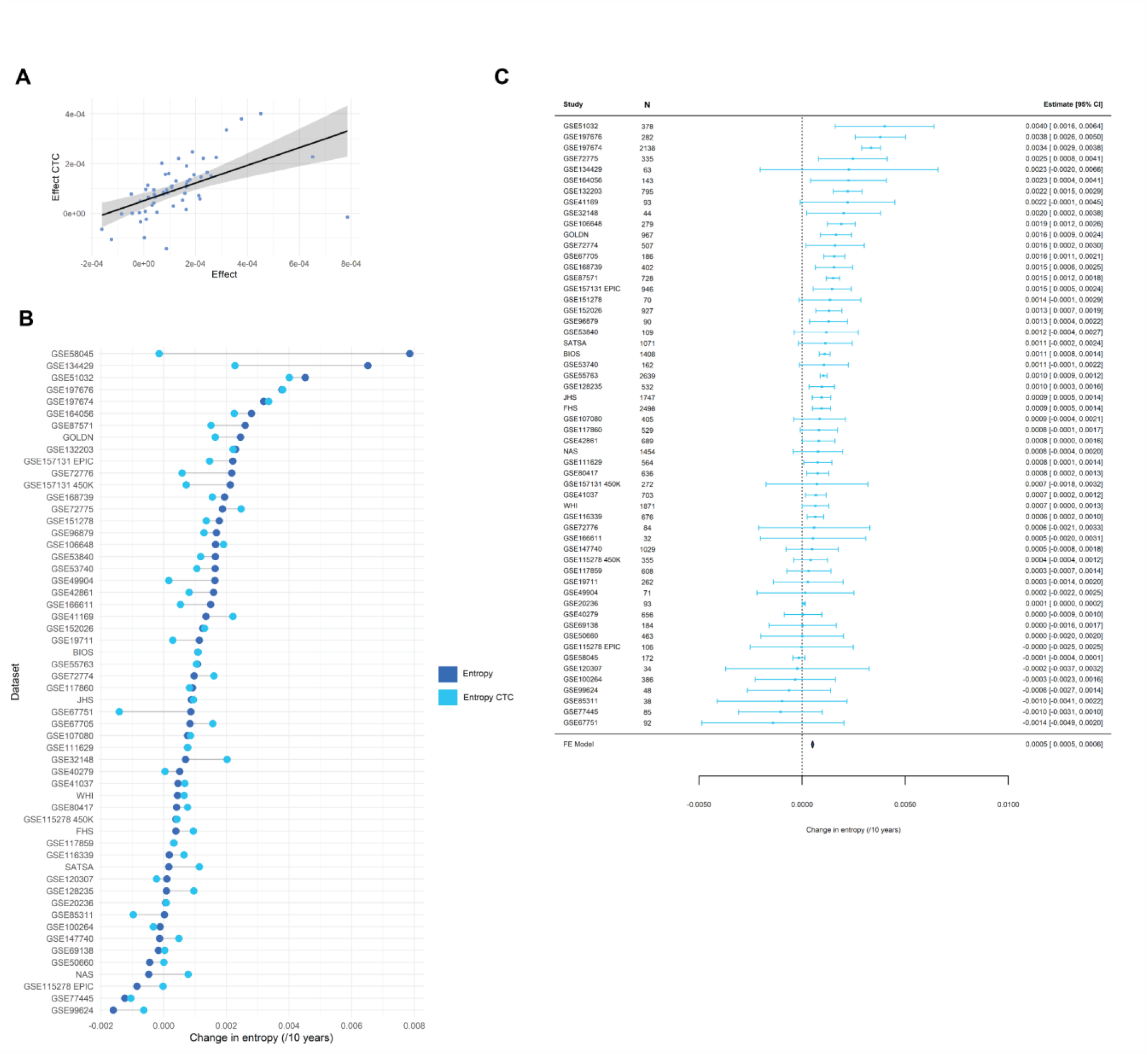
Entropy changes in blood after correction for cell types. **A)** The effect sizes from the independent EWAS of entropy and age for the analyses unadjusted for the 5 largest blood cell types (x-axis) and the effect size for the cell type corrected (CTC) analyses. Each point on the graph represents a single blood dataset. The Pearson’s correlation is 0.53. **B)** A lollipop chart to display the change in direction of entropy before (royal blue) and after cell type correction (skyblue) for each dataset (y-axis). **C)**A forest plot of the genome-wide meta-analysis of entropy and age in the cell type corrected datasets. The change in entropy per decade of age is located on the x-is, with the effect size and standard error measurements from the independent EWAS on the y-axis. The 95% confidence intervals are on the right pane. The meta-analysis effect size is represented by the navy polygon at the bottom of the plot.

**Extended Data Fig. 9.**
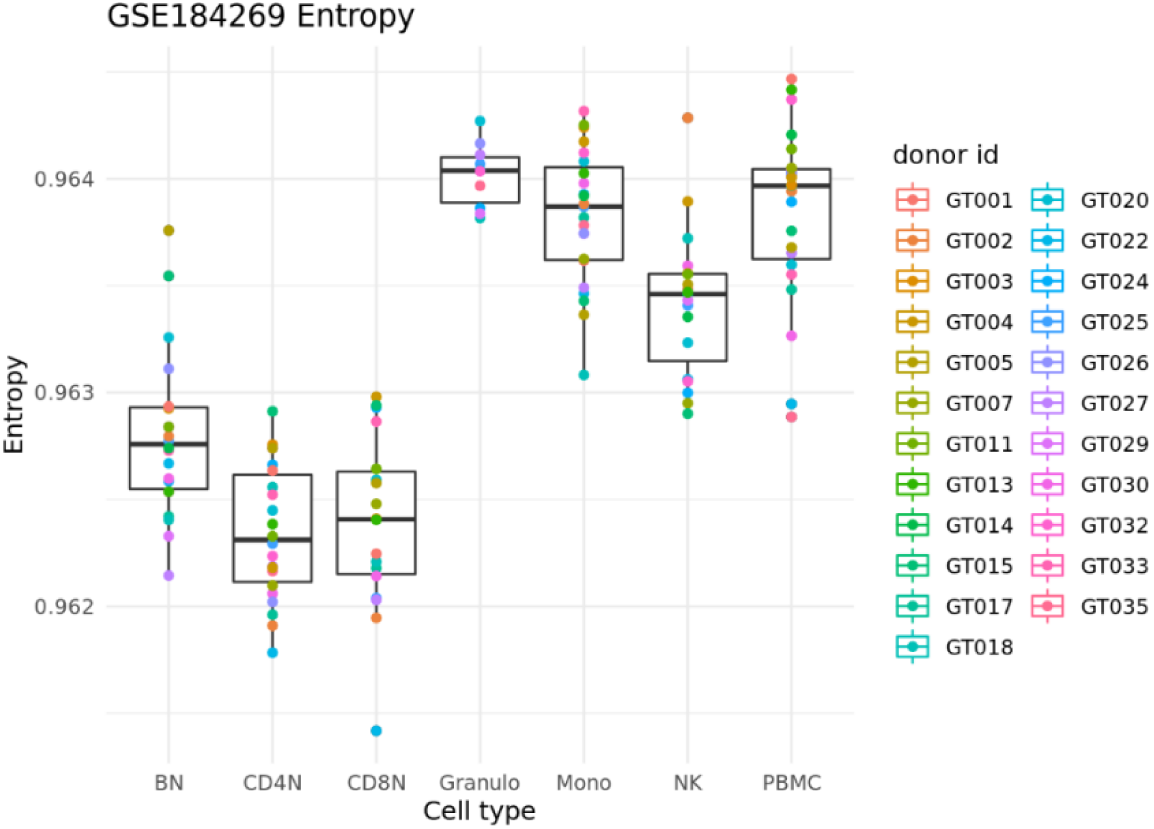
Entropy comparison in six sorted blood cell types in a single dataset. Individual entropy values (y-axis) for six sorted cell types (naïve B cells, naïve CD4+ T cells, naïve CD8+ T cells, granulocytes, monocytes, natural killer cells) and peripheral blood mononuclear cells (PMBCs) (x-axis) from the GSE184269 dataset that contains both mixed and sorted blood cells from the same individuals. Each individual is colour-coded according to the legend (donor id) on the right side of the chart.

**Supplementary Table 1.**
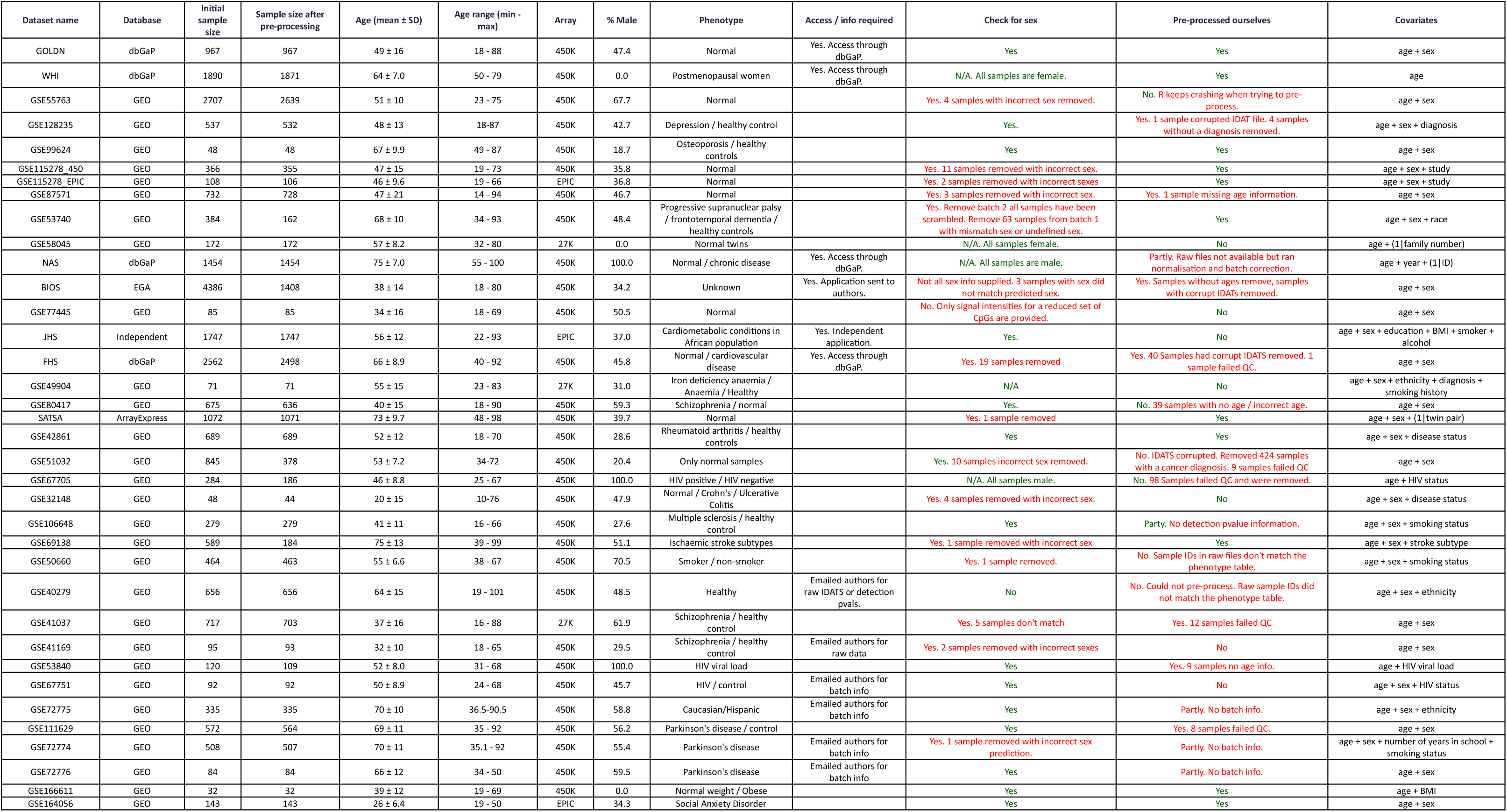
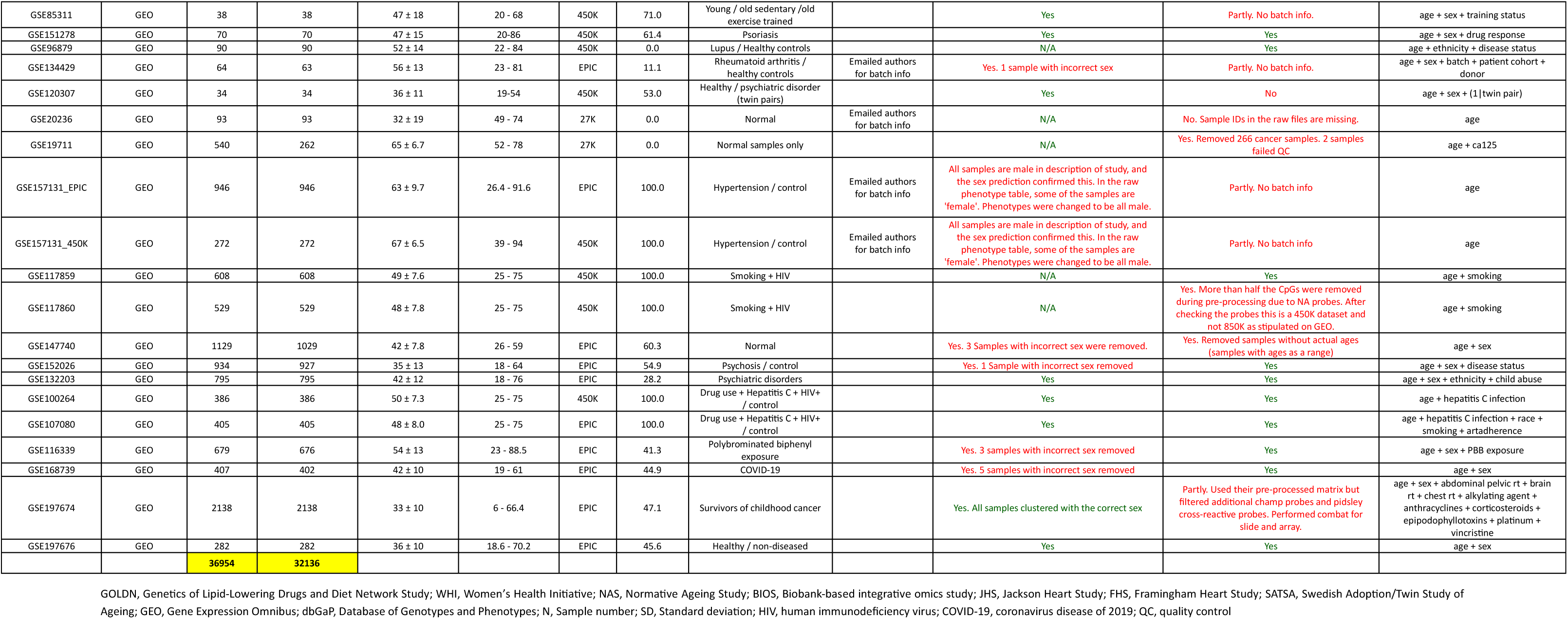
Description of blood datasets.

**Supplementary Table 2.**
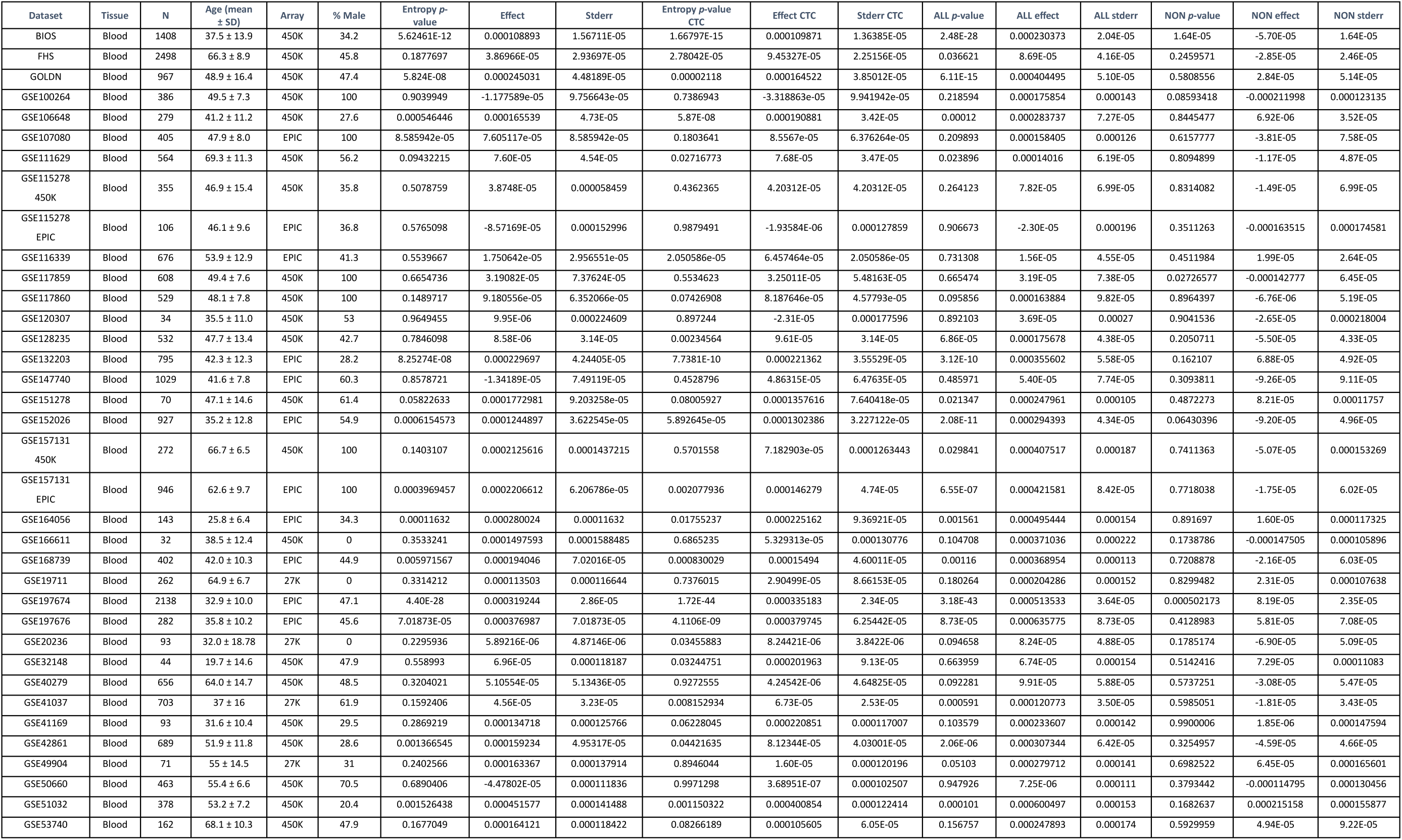
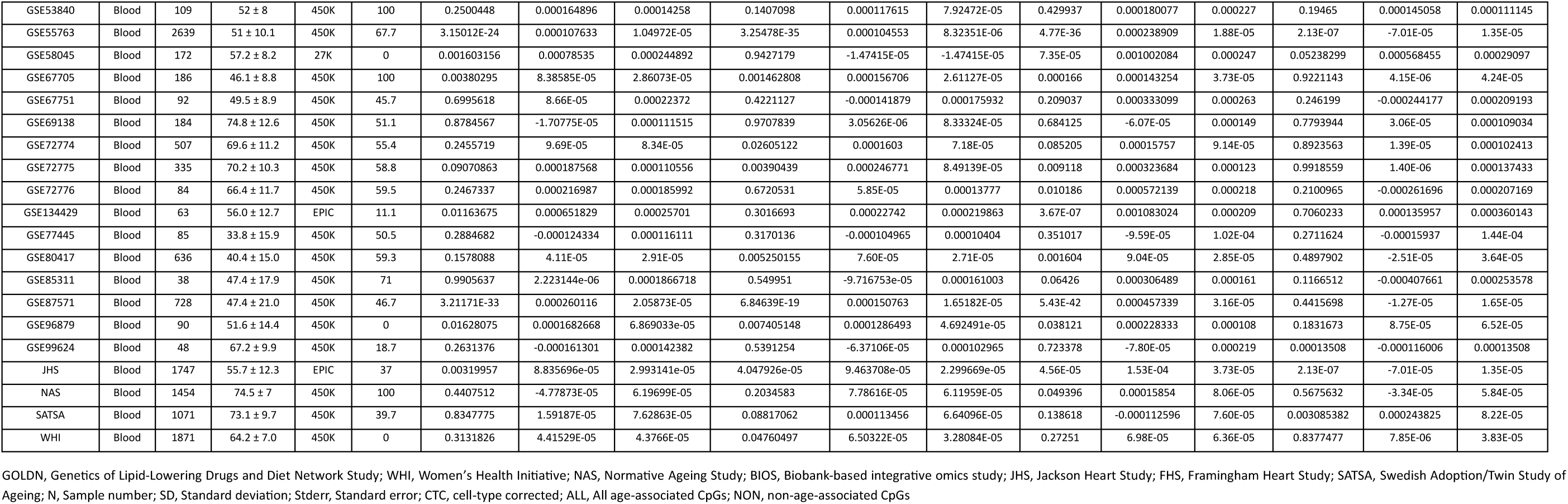
Shannon entropy results in blood.

**Supplementary Table 3.** Entropy results for three categories of age-associated CpGs.

| A | DMPs only |  |  | DMPs only CTC |  |  |
| --- | --- | --- | --- | --- | --- | --- |
|  | p-value | effect | stderr | p-value | effect | stderr |
| BIOS | 0.00263494 | 0.000046 | 1.53E-05 | 0.001221366 | 0.000046509 | 1.44E-05 |
| GSE87571 | 2.13E-18 | 0.000176 | 1.96E-05 | 2.85E-10 | 0.000111197 | 1.74E-05 |

| B | DMP/ VMP |  |  | DMP / VMP CTC |  |  |
| --- | --- | --- | --- | --- | --- | --- |
|  | p-value | effect | stderr | p-value | effect | stderr |
| BIOS | 1.29E-41 | 0.000383 | 2.75E-05 | 5.30E-78 | 0.0003835 | 1.92E-05 |
| GSE87571 | 1.88E-47 | 0.000649 | 4.16E-05 | 2.81E-32 | 0.000352149 | 2.83E-05 |

| C | VMPs only |  |  | VMPs only CTC |  |  |
| --- | --- | --- | --- | --- | --- | --- |
|  | p-value | effect | stderr | p-value | effect | stderr |
| BIOS | 1.18E-12 | -0.0000900 | 1.25E-05 | 3.84E-11 | 0.00030754 | 3.08E-04 |
| GSE87571 | 0.02844887 | -0.0000415 | 1.89E-05 | 0.03538065 | -0.00003671 | 1.74E-05 |

BIOS, Biobank-based integrative omics study; DMPs, differentially methylated positions; VMPs, variably methylated positions; stderr, standard error; CTC, cell type correction.

**Supplementary Table 4.**
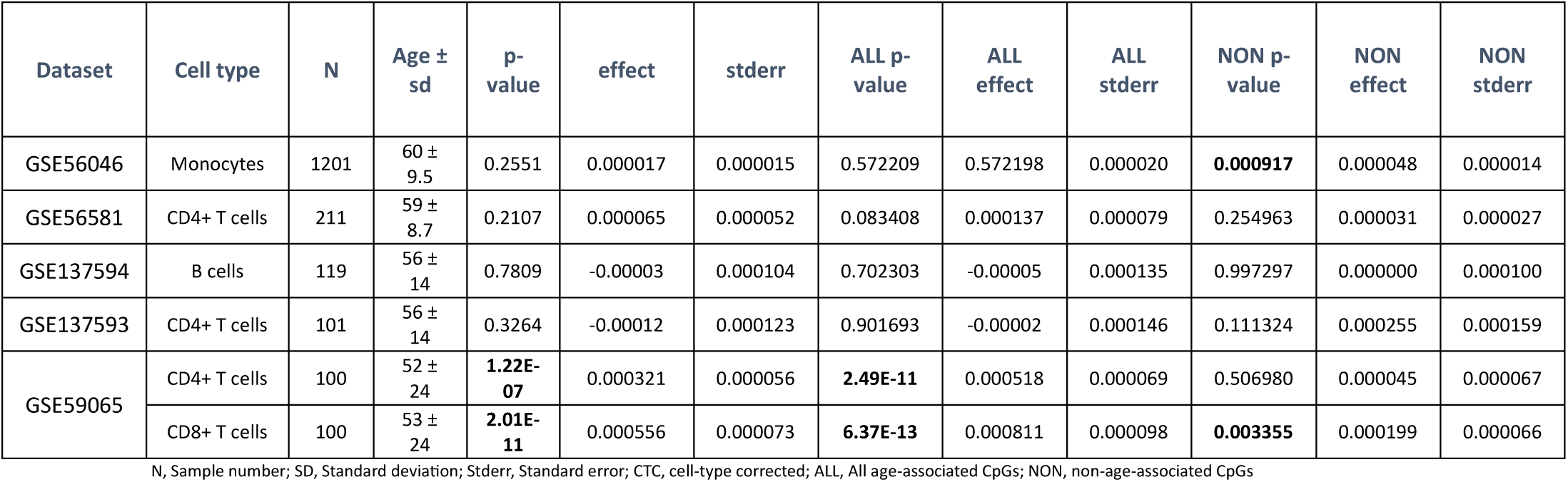
Entropy results in datasets containing isolated cell types.

